# A hybrid model combining evolutionary probability and machine learning leverages data-driven protein engineering

**DOI:** 10.1101/2022.06.07.495081

**Authors:** Alexander-Maurice Illig, Niklas E. Siedhoff, Ulrich Schwaneberg, Mehdi D. Davari

**Affiliations:** Institute of Biotechnology, RWTH Aachen University, Worringerweg 3, Aachen, 52074, North Rhine-Westphalia, Germany; DWI-Leibniz Institute for Interactive Materials, Forckenbeckstraße 50, Aachen, 52074, North Rhine-Westphalia, Germany; Department of Bioorganic Chemistry, Leibniz Institute of Plant Biochemistry, Weinberg 3, Halle, 06120, Saxony-Anhalt, Germany

**Keywords:** Protein engineering, machine learning, protein sequence encoding, direct coupling analysis, hybrid modeling

## Abstract

Protein engineering through directed evolution and (semi-)rational approaches has been applied successfully to optimize protein properties for broad applications in molecular biology, biotechnology, and biomedicine. The potential of protein engineering is not yet fully realized due to the limited screening throughput hampering the efficient exploration of the vast protein sequence space. Data-driven strategies have emerged as a powerful tool to leverage protein engineering by providing a model of the sequence-fitness landscape that can exhaustively be explored *in silico* and capitalize on the high diversity potential offered by nature However, as both the quality and quantity of the inputted data determine the success of such approaches, the applicability of data-driven strategies is often limited due to sparse data. Here, we present a hybrid model that combines direct coupling analysis and machine learning techniques to enable data-driven protein engineering when only few labeled sequences are available. Our method achieves high performance in predicting a protein’s fitness based on its sequence regardless of the number of sequences-fitness pairs in the training dataset. Besides reducing the computational effort compared to state-of-the-art methods, it outperforms them for sparse data situations, i.e., **50 − 250** labeled sequences available for training. In essence, the developed method is auspicious for data-driven protein engineering, especially for protein engineers who have only access to a limited amount of data for sequence-fitness landscape modeling.

## 1 Introduction

The natural evolution of organisms is driven by random genotypic mutation spawning proteins adapted for numerous specific tasks. The ability to engineer proteins and thereby overcome certain *ex vivo* limitations, e.g., expression, binding, activity, thermostability, or promiscuity [1], has made proteins accessible for broad applications in molecular biology, biotechnology, and biomedicine [2–5]. For protein engineering, strategies based on directed evolution [6, 7], (semi-)rational methods [8, 9], and approaches combining directed evolution and computational analysis such as KnowVolution [10] are applied, offering almost unlimited possibilities for tailoring proteins.

Modern technologies such as high-throughput screening (HTS), next generation sequencing, and deep mutational scanning (DMS) allow to sequence and screen protein variants in an automated fashion, consequently generating huge protein sequence databases [11]. However, establishing such HTS systems is very elaborate such that they are not implemented in most protein engineering campaigns [12].

Despite significant progress in recent years in the areas of high-throughput screening and DNA sequencing [13], our ability to realize the full potential of protein engineering remains severely limited in the absence of HTS systems, as it hampers the identification of beneficial positions and results in a lack of information required for beneficial recombination. Therefore, most experimental methods explore only a small subspace of the vast sequence space leading to a local maximum, when interpreting protein engineering as an optimization problem.

In recent years, machine learning (ML) and statistical approaches have expanded the toolbox of protein engineers by extracting knowledge from data for constructing a model of the protein sequence-fitness landscape that can subsequently be explored *in silico* [11, 14]. Such a model not only enables *in silico* screening, which has orders of magnitude higher throughput compared to experimental methods [15] while minimizing costs, but can also support exploration and recombination. ML methods applied for predicting a protein’s fitness from its sequence are mostly supervised and use numerical representations of protein sequences as features (independent variables) and the corresponding observed fitness (assay-based dependent variables) as labels [12, 16]. A fundamental task in the construction of such models is to map the primary sequence to a representation that provides as much information as possible about the sequence necessary for fitness prediction [14]. Hence, diverse strategies have been developed for encoding that can mainly be divided into amino acid-specific [15, 17–21] and sequence-specific encoding [22–25]. Various studies show that representations tailored to the sequence of interest usually lead to better models than those that make use of more general amino acid properties for encoding [23, 26]. As a result, frameworks integrating such tailored encoding techniques have been developed, e.g, eUniRep [23] and ECNet [24]. Multiple sequence alignments (MSAs) serve as a fruitful data source for generating such customized representations [27, 28], which is often achieved by training recurrent neural networks, variational autoencoder, or transformer architectures in an unsupervised manner on the MSA [14, 22, 23, 29] or inferring probabilistic models, e.g., (generalized) Potts-Models [30]. In contrast to supervised machine learning, such statistical methods considering direct couplings, e.g. EVmutation [31], were developed to predict protein-protein interactions [32, 33] or protein structures [34, 35] and allow to generate an output that strongly correlates with the fitness of the sequence under investigation in the absence of labeled data [31, 36]. Recently Luo et al. and Hsu et al. incorporated direct coupling analysis (DCA)-derived terms for constructing machine learning models [24, 26]. However, as supervised ML methods require minimum amount of sequence-fitness pairs to generalize well [22], the performance of unsupervised methods is limited and cannot be tuned within protein engineering campaigns, which are often designed iteratively. Furthermore, sequence-specific encoding functions usually produce high dimensional representations enhancing the risk of over-fitting for sparse training datasets.

In order to overcome these limitations, we herein present an approach that combines a machine learning model trained on sequence-fitness pairs with the statistical energy [31] of a DCA model. Further, we introduce a new encoding strategy allowing to generate low-dimensional sequence-specific representations, minimizing the risk of overfitting. Testing our approach on 19 single-saturated and 2 DMS engineering datasets (Supplementary Tab. 1) revealed, that such a hybrid model represents a computationally efficient and highperforming strategy for low-to high-*N* protein engineering tasks. In particular, our method raises the prospect of building trustworthy models even with sparse data.

## 2 Results

To generate a model achieving high performances for various training datasets – from sparse to rich –, we developed a hybrid model by combining a linear model trained on sequences with experimentally measured labels with a probability feature derived from evolutionary related sequences. The components of our hybrid model are both based on evolutionary features and were derived applying DCA on MSAs constructed by sequence similarity searches. An overview of the entire workflow for generating such a hybrid model is depicted in Fig. 1.

**Fig. 1:**
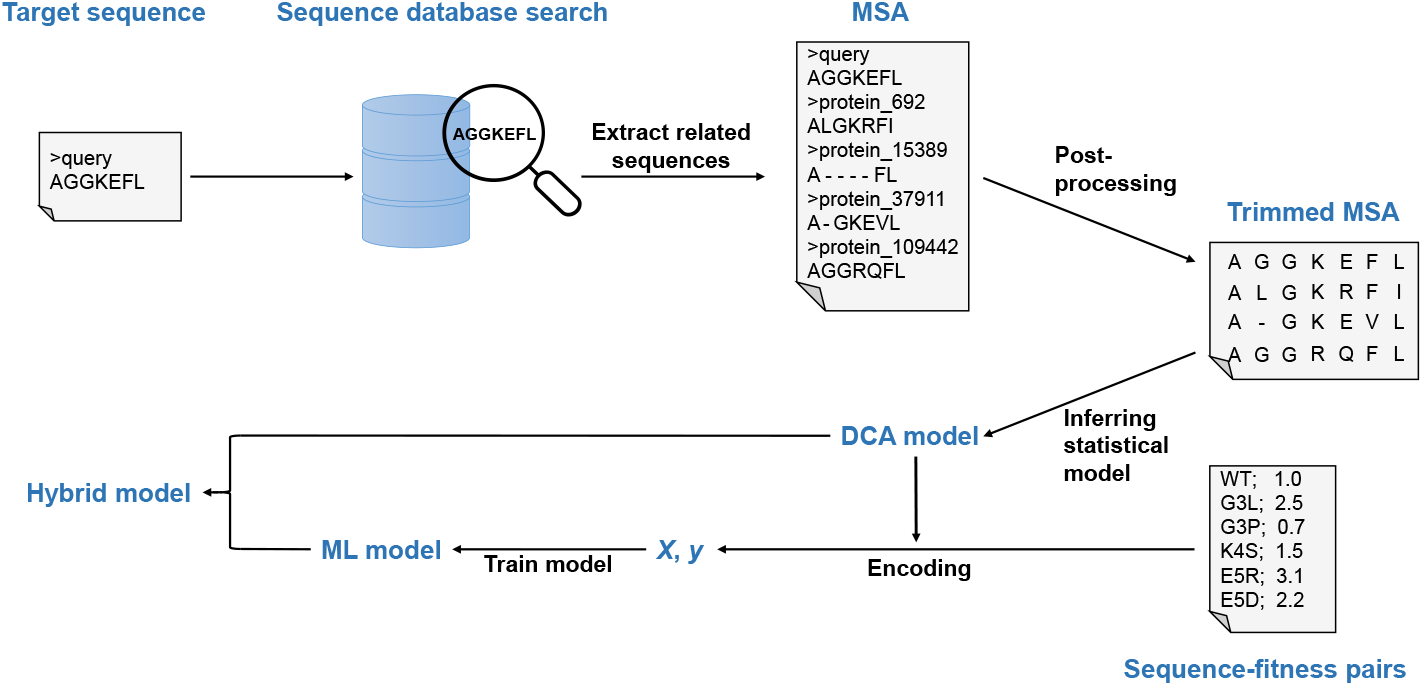
Workflow for constructing a hybrid model for predicting protein fitness from sequence. First, a homology search is performed to construct an MSA, which is subsequently post-processed resulting in a trimmed and curated MSA. In the next step, a direct coupling analysis (DCA) model is inferred and used to predict the statistical energy of a sequence. Further, its parameters are utilized for encoding the labeled sequences ***X*** before subjecting them and the associated fitness values ***y*** to the ML model training. Both models’ predictions are blend and merged to construct the hybrid model.

We compared our approach in terms of model performance to other state-of-the-art methods that also use evolutionary encoding techniques to build ML models and EVmutation. Besides, we tested a machine learning model based on a simple form of encoding as a baseline approach and a pure (non-hybrid) machine learning model based on our proposed encoding. Details of the preprocessing steps for feature generation, hybrid model implementation, and the implementation of all other methods tested, can be found in Section 4. More detailed information about the algorithmic flow of our hybrid model is provided in the Supplementary Information (Supplementary Fig. 1).

### 2.1 Comparison of Performance of Hybrid Model with State-of-the-art Methods

To analyze our approach’s applicability as a tool for accelerating protein engineering campaigns, we studied its performance in terms of Spearman’s *ρ* on 19 public datasets. We compared it to state-of-the-art methods, i.e., ECNet [24], eUniRep [23], EVmutation [31], and UniRep [22]. We used Spearman’s *ρ* as metric of model performance because we consider the overall ranking of variants to be more relevant for protein engineers than predicting the fitness of unknown variants with the highest accuracy. Besides, Spearman rank correlation coefficient was used to allow comparison between different methods, i.e. the statistical and supervised models.

As a baseline we chose a one-hot encoded linear model. These 19 datasets consist of data on single substituted variants (variant and fitness labels). They are hereafter referred to as single-saturation mutagenesis (SSM) datasets, although they do not necessarily cover the full natural diversity (also see Hopf et al. [31]). For a detailed description of the datasets and methods used, the reader is referred to Sec. 4.

In our study, we defined the protein engineering task as ranking the hold-out variants correctly according to their experimentally determined label. For estimating the suitability of the different methods for varying data situations, e.g., from sparse to fully diverse, we tested the model performance for the protein engineering task for models trained on a low to a high number of variants. To compare EVmutation with the supervised models, the performance was estimated on the differently sized holdout sets. The given performance *ρ* for a specific size of the training dataset of a model represents the arithmetic mean of the individual performances achieved on ten different splits. To ensure comparability, the same splits were used to evaluate each model.

Besides, we studied the ability of substitutional extrapolation on two datasets by training on single substituted variants and predicting higher substituted variants, i.e., ranking associated (i) enrichment scores of *Saccharomyces cerevisiae* poly-A binding protein (PABP) and (ii) brightness of *Aequorea victoria* green fluorescent protein (avGFP) variants.

#### 2.1.1 Hybrid Model Achieves High Performance Independent of the Size of the Training Dataset

To study the dependency of the model performance on the size of the training dataset, we successively increased the training dataset from *N*_train_ = 15 up to *N*_train_ = 1000. We evaluated the ranking of the remaining variants in the dataset. For datasets comprising less than 1000 variants, we limited the size of the training dataset to 80 %. In addition to the fixed absolute sizes for training, we determined the performance of each model when trained on 80 % of all data. To derive general trends, we present the averaged model performance on the 19 datasets

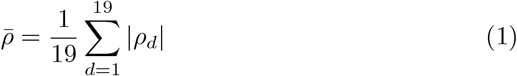

for each approach hereinafter. However, the individual case studies are provided in the Supplementary Information (Supplementary Fig. 2).

**Fig. 2:**
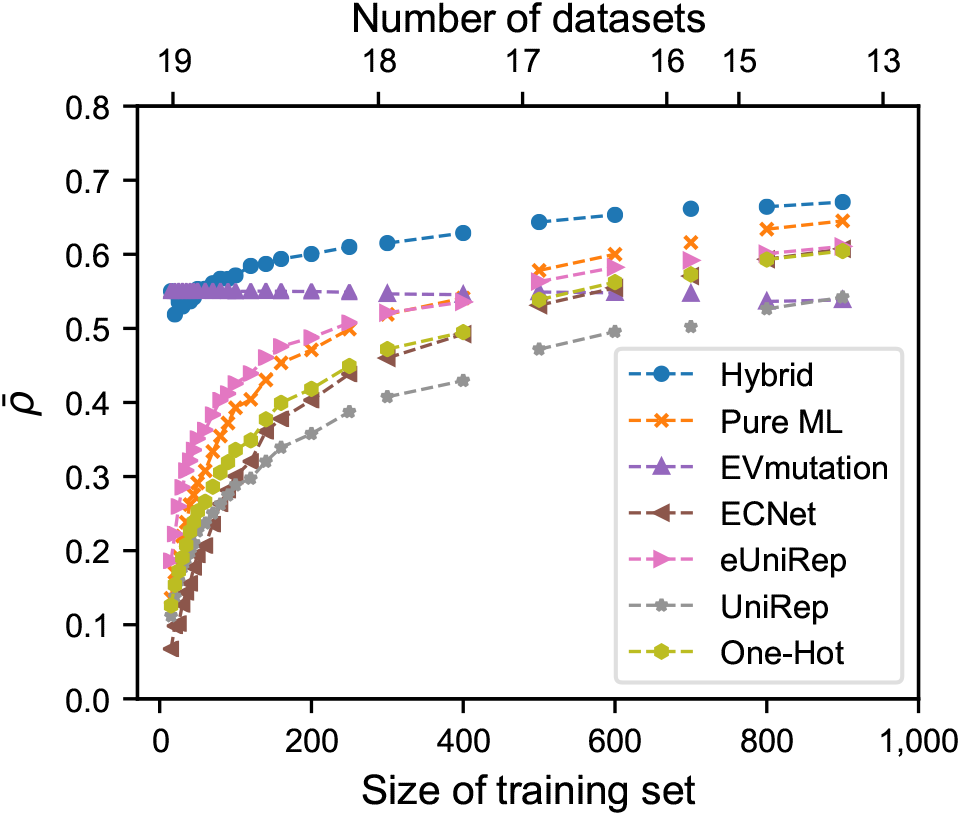
Averaged performance of different methods for ranking the variants according to their fitness label at different training dataset sizes. The given performance metric 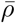 (averaged for the datasets and the 10 splits tested) quantifies the ability to rank the holdout variants correctly according to their labeled fitness. The performance is estimated only if at least 20 % of the entire dataset remains for prediction. Thus, as the size of the training dataset increases, the total number of datasets available for determining the model performance decreases.

As can be seen in Fig. 2, the hybrid model performed best on average when trained on up to 1000 variants. However, for sparse data regions, with *N*_train_ in the range of 15 to 50, EVmutation turned out to be superior. Furthermore, while EVmutation did not depend on the size of the training dataset, the performance of all supervised methods did strongly increase with increasing training data. On average, the hybrid model outperformed EVmutation at sizes of the training datasets of ≥ 50. In contrast to the hybrid model, the other machine learning methods did not show high performances for *N*_train_ ≤ 250. While the pure ML approach, eUniRep, one-hot, and ECNet outperformed EVmutation on average at sizes of the training datasets of 500 to 600, UniRep reached similar performances only at sizes of 800 to 900. With a size of the training dataset of 1000, the hybrid model ranks best, followed by the pure ML model and the similarly performing models one-hot, ECNet, and eUniRep, finally followed by EVmutation and UniRep.

In addition to studying the performance as a function of the size of the training dataset, models trained on 80 % of the available data were generated and evaluated based on their ability to correctly predict the fitness rank of the variants in the remaining 20 % of the data. Here, we found that while the hybrid model still performed best on average across all datasets, the pure ML model, eUniRep, and ECNet performed similarly well, closely followed by UniRep and finally EVmutation (Tab. 1). ECNet and the hybrid model performed equally well regarding the median performance (*ρ* = 0.71). Furthermore, it turned out that all supervised models have a minimum performance of approximately 0.40, while EVmutation falls significantly below this value. A close look at the maximum achievable performance on the tested datasets shows that all methods can reach high Spearman’s *ρ* of at least 0.75.

Fig. 3 illustrates the performances achieved for each model by presenting the mean absolute Spearman rank coefficient of correlation of ten different splits |*ρ*| obtained for each of the 19 datasets when trained on 80 % of the data and tested on the remaining data. The individual standard errors on the means are not shown because they are marginal and displaying them would affect the clarity. As can be seen in Fig. 3, there is no one method that is superior for prediction tasks of all datasets. However, the hybrid model can achieve the highest performance for at least 9 of the 19 datasets studied. In contrast, EVmutation performs on 13 datasets the worst. Since the hybrid model contains this approach to a certain extent, the findings highlight the impact of supervised training on performance. The superiority of the hybrid model is surprising given the small number of parameters compared to the other encoding techniques. However, it is to be expected that models with many parameters, e.g., ECNet, will likely outperform the hybrid model for large training datasets. We also tested nonlinear models and compared the obtained performances with the performances of the linear models. The comparison of the four different supervised pure ML models for this engineering task, i.e., Ridge, Lasso, support vector, and random forest regression trained on 80 % of the data, is provided in the Supplementary Information (Supplementary Fig. 3 and Supplementary Tab. 2). It is noteworthy that nonlinear models did not drastically improve model performance. Only support vector machine-based regression showed slightly improved performance for this engineering task than both linear models, whereas random forest-based regression showed lower performance. Due to the increased risk of overfitting the non-linear model and the similar performances achieved, we retain linear Ridge regression as the default method for modeling.

**Table 1:** Comparison of the hybrid model with different methods when ranking the variants according to their fitness label for all SSM datasets. Mean, median, minimum, and maximum absolute Spearman rank correlation coefficients observed on 19 different SSM datasets for the models under investigation when trained and tested on 80 % and 20 % of the data, respectively. The errors on the median, minimum, and maximum value correspond to the standard error of the mean obtained from 10 different splits for the associated dataset.

| Model | Mean | Median | Minimum | Maximum |
| --- | --- | --- | --- | --- |
| Hybrid | 0.69 | $0.71 \pm 0.03$ | $0.45 \pm 0.05$ | $0.86 \pm 0.01$ |
| Pure ML | 0.68 | $0.70 \pm 0.02$ | $0.46 \pm 0.03$ | $0.85 \pm 0.01$ |
| EVmutation | 0.55 | $0.55 \pm 0.04$ | $0.23 \pm 0.06$ | $0.75 \pm 0.01$ |
| ECNet | 0.66 | $0.71 \pm 0.02$ | $0.41 \pm 0.08$ | $0.83 \pm 0.02$ |
| eUniRep | 0.67 | $0.70 \pm 0.04$ | $0.45 \pm 0.03$ | $0.82 \pm 0.01$ |
| UniRep | 0.62 | $0.65 \pm 0.02$ | $0.41 \pm 0.08$ | $0.81 \pm 0.03$ |
| One-Hot | 0.64 | $0.70 \pm 0.03$ | $0.41 \pm 0.09$ | $0.76 \pm 0.02$ |

**Fig. 3:**
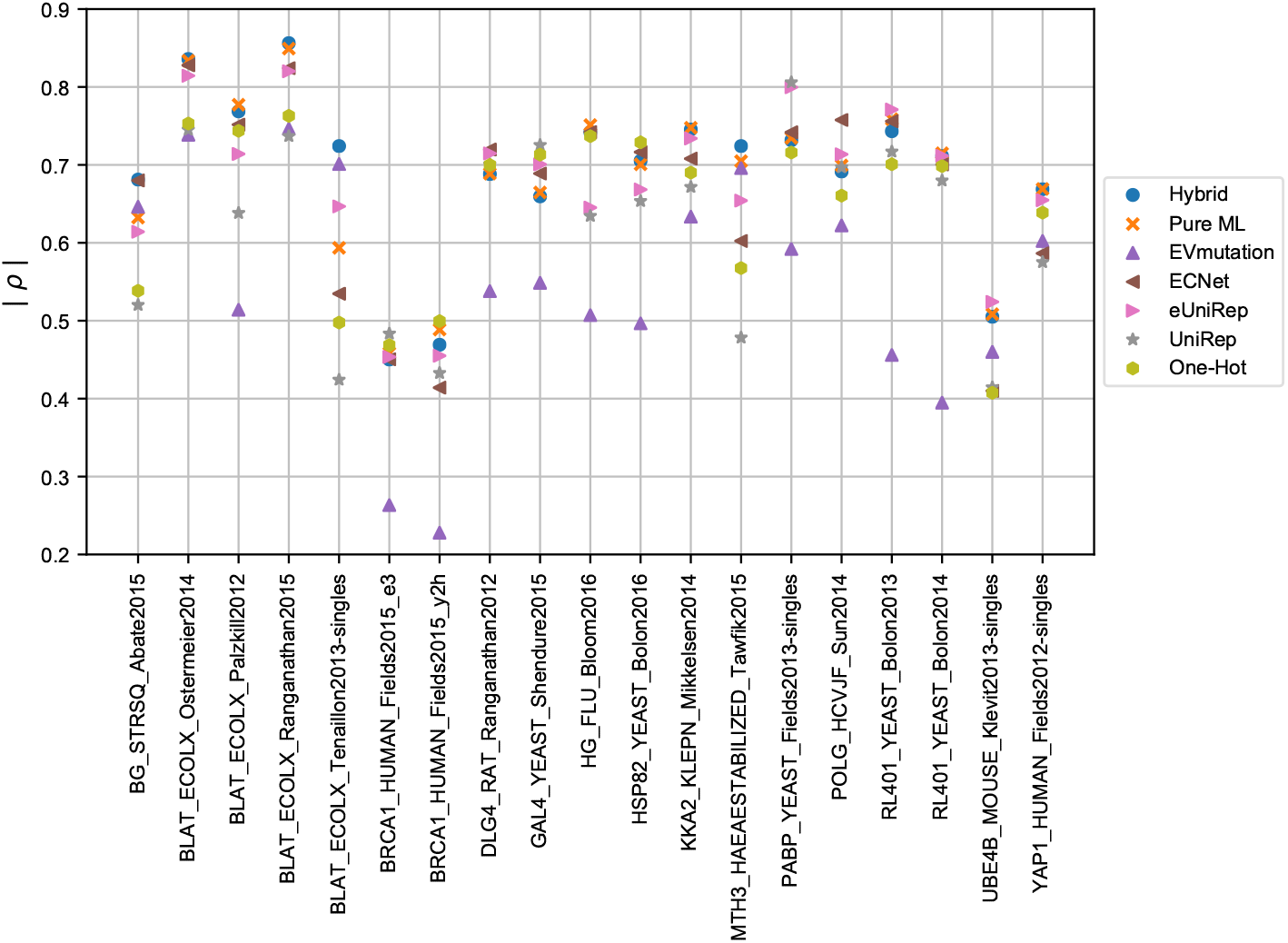
Performances of different methods for ranking the variants according to their fitness label for all SSM datasets. Absolute Spearman’s rank coefficient of correlation |*ρ*| achieved for each of the 19 SSM datasets when trained on 80 % of the data and tested on the remaining 20 %. The given performance represents the mean of ten different splits. Since the errors on the mean are very small, they are not shown in the figure for clarity.

#### 2.1.2 Supervised Model Gains Weight with Increasing Training Dataset

The hybrid model combines the output of a statistical model with the output of a linear model constructed by supervised training. The contribution of the individual terms is controlled by the parameters *β*_1_ and *β*_2_, respectively, and was examined at different sizes of the training dataset. Fig. 4 illustrates the mean values of the parameters *β*_1_ and *β*_2_ for all datasets. As can be seen, the influence of the supervised model increases from *β*_2_ = 0.0 up to approximately 0.7 with the increasing size of the training dataset, whereas the mean of *β*_1_ decreases from 1.0 to almost 0.0. This behavior is to be expected since, as previously shown, the performance of supervised models increases as the training dataset increases. The contributions of both *β*_1_ as well as *β*_2_ at the different training dataset sizes for all 19 SSM datasets can be obtained from the Supplementary Information (Supplementary Fig. 4).

**Fig. 4:**
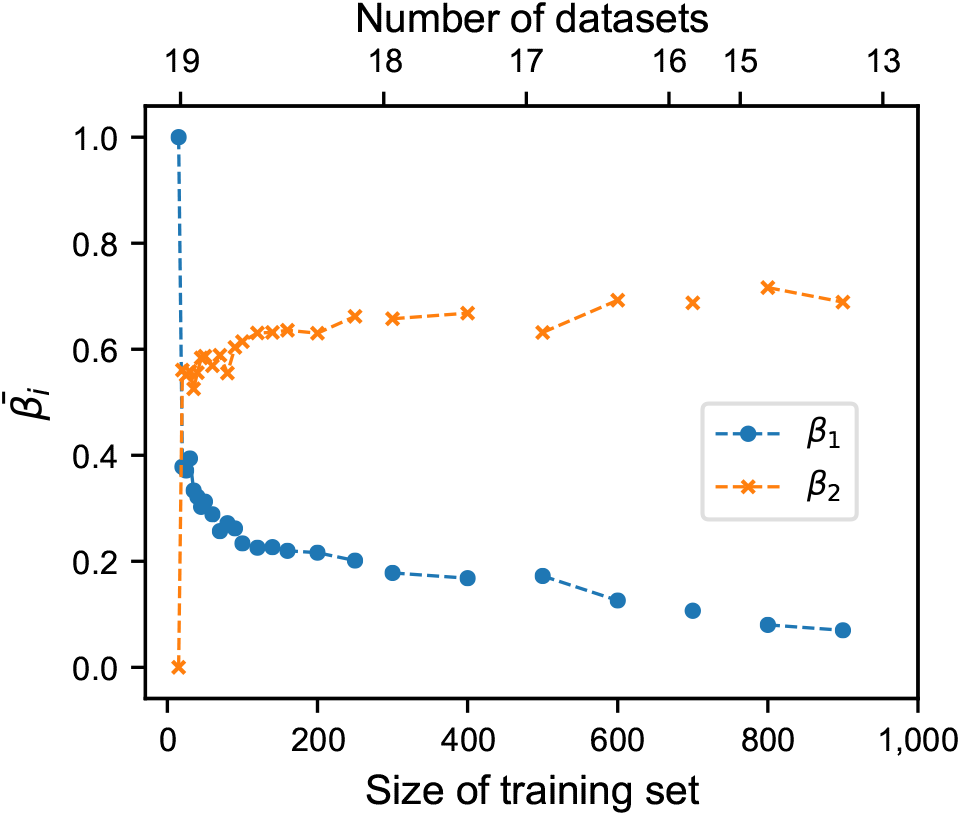
Influence of the DCA and ML model on the predicted fitness of the hybrid model at different training dataset sizes. The hybrid model results after combining both models with their individual weights, *β*_1_ (DCA model contribution) and *β*_2_ (ML model contribution), and is constructed only if at least 20 % of the full dataset remains for prediction.

#### 2.1.3 Hybrid Model Allows Prediction of Improved Recombinants Already with Small Training Datasets

We studied the ability of the hybrid model for extrapolation on two datasets by training on single substituted variants and predicting higher substituted variants, i.e., ranking (i) the enrichment scores of PABP variants and (ii) the brightness of avGFP variants. In addition, extrapolation studies were performed for the pure ML, one-hot model, the EVmutation approach, and a simple additive model for comparison. Since the additive model generates the fitness predictions of multi-substituted variants by summing up the measured labels of the individual single-substituted variants it contains (Sec. 4), this approach is only suitable for recombining known variants. Similar to the previous performance studies, the ability to correctly rank multi-substituted variants was investigated as a function of the size of the training dataset. It should be noted that for all supervised models *ρ* represents the mean of 10 different splits.

##### Model Performance for Substitutional Extrapolation Task of Poly-A Binding Protein

The PABP dataset consists of 1, 142 sequences possessing one amino acid substitution and 33, 771 double substituted variants [37]. As depicted in Fig. 5 – and elucidated in the previous analyses, the hybrid model is superior to the other methods in regions with 50 < *N*_train_ < 500. For all supervised models, an increase in performance is observed as the size of the training dataset increases. While the performance for the hybrid and the pure ML strategy stagnates for larger training datasets (*N*_train_ ≥ 500) at a value of approximately *ρ* = 0.75, outperforming the probability density approach EVmutation, which reaches *ρ* = 0.65, the performance of the one-hot model increases continuously up to *ρ* = 0.85. Interestingly, the pure ML as well as the one-hot encoded model share an almost identical behavior for 20 ≤ *N*_train_ ≤ 400. The further increase in the performance of the one-hot model illustrates the frequent problem of overfitting to the training subset used for validation [38, 39], since it can provide one independent parameter to each variant. Since the regularization parameter adjusts to the smallest possible value of 10^−6^ of the grid after a fivefold cross validation on all single variants, the model is almost nonregularized and thus transitions to the additive model. It is expected that for datasets with dominant epistatic effects, the additive model will perform significantly worse, as it can not capture such non-linearities. The same applies to the one-hot model, when trained on single-substituted variants only.

**Fig. 5:**
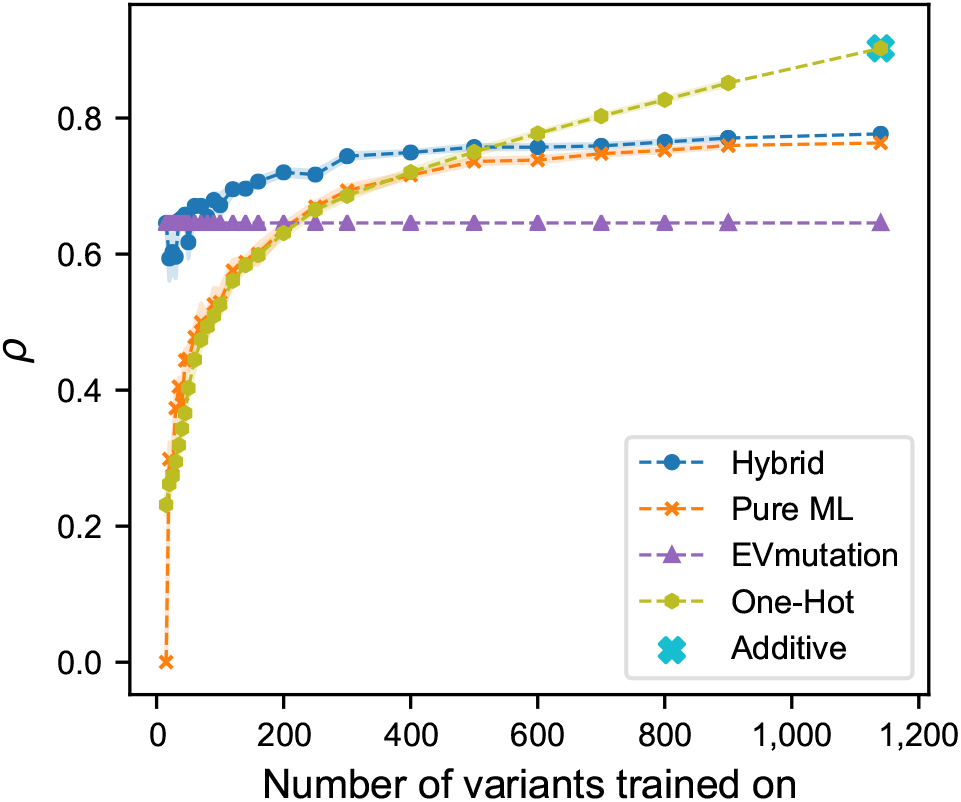
Performance of different methods for ranking recombinants of the poly A-binding protein according to their fitness label. Spearman’s rank coefficient of correlation *ρ* was determined after ranking double substituted variants of the poly-A binding protein as a function of the number of single substituted variants in the training dataset. The colored area represents the standard error of the mean.

##### Model Performance for Substitutional Extrapolation Task of Green Fluorescent Protein

To study the ability of extrapolation for avGFP, we followed the same procedure as PABP. However, as the avGFP dataset represents the outcome of a deep mutational scan with multiple mutations combined, it allows for studying the ability to correctly rank higher substituted variants ranging from 2 to 6 (Fig. 6 and Supplementary Fig. 5), when trained on single substituted variants.

**Fig. 6:**
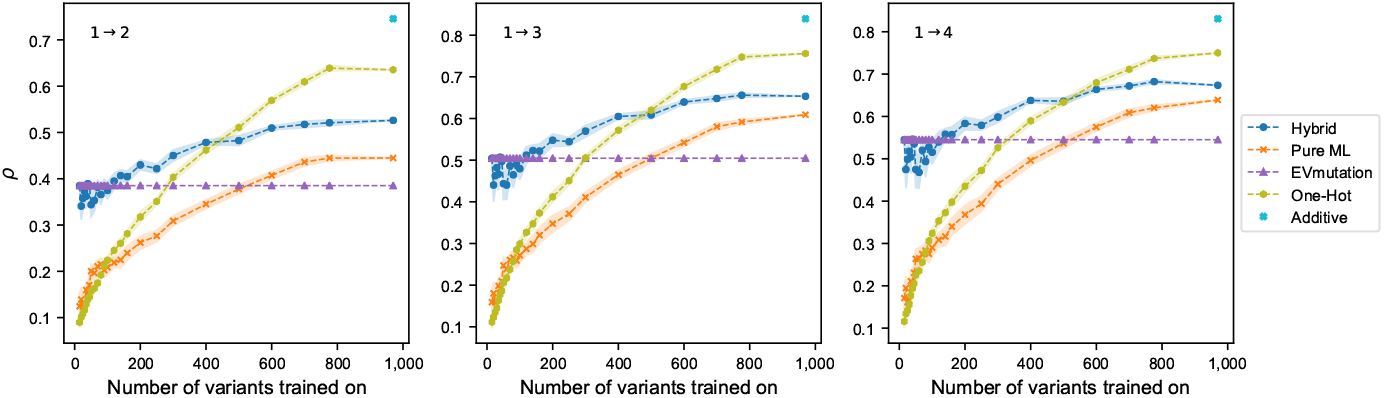
Performance of different methods for ranking recombinants of the green fluorescent protein according to their fitness label. Spearman’s rank coefficient of correlation *ρ* was determined after ranking multiple substituted variants with two (1 →2), three (1 →3), or four (1 → 4) simultaneous substitutions of the green fluorescent protein from *Aequorea victoria* as a function of the number of single substituted variants in the training dataset. The colored area represents the standard error of the mean.

It can be seen from Fig. 6 that the overall performances are significantly lower than in the PABP extrapolation task. Nevertheless, it can be observed that the hybrid model is superior in the range of 50 < *N*_train_ < 500. Although the pure ML model outperforms the EVmutation approach at *N*_train_ ≥ 500, it is no longer consistent with the one-hot model. After training on all single substituted variants, the additive model performs best, followed by the one-hot, hybrid, pure ML, and EVmutation approach. This behavior can be explained by the number of available parameters as described before. Interestingly, however, the one-hot model cannot reach the performance of the additive model, which is due to the *L*^2^-regularization. As described in more detail in Sec. 4, predicting the fitness of unknown substitutions at a specific position corresponds to returning the influence of all known substitutions at this position for a regularized one-hot model. While this assumption holds for the PABP dataset, i.e., it shows that the summed fitness values of a position are linearly related to the summed coefficients at this position, it is invalid for the avGFP dataset. As a result, a significantly increased regularization and lowered performance is observed compared to the additive model. Even though the additive model achieved the best ranking for the case studies, it is important to mention that this model is limited to recombination and cannot account for epistatic effects. In addition, it was shown that the assumptions interwoven in the regularized one-hot model do not hold. Such a model is massively affected by overfitting in contrast to the other models (see train and test performances in the Supplementary Tab. 3 and Supplementary Tabs. 4 - 7).

### 2.2 Curating the MSA is Crucial for Model Performance

As the MSA can contain positions with high gap content, it is necessary to exclude these positions when inferring the statistical model, to avoid bias. Often, sequences within an MSA share the same evolutionary history. Within this picture, frequently occurring characters at a specific position can be interpreted as highly conserved amino acids. Since the probabilistic model is – among others – constrained to the frequency

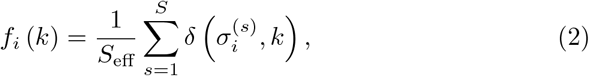

with *S* being the number of total sequences and *S*_eff_ being *S* reduced by the number of gaps at position *i* in the MSA, *k* a representative of one of the 20 proteinogenic amino acids, and 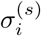 the amino acid representative at position *i* of the *s*-th sequence in the MSA, changing *f*_*i*_ will influence Δ*E* [33].

Considering Eq. 2, an increasing gap content at a specific position leads to an increase of *f*_*i*_ that causes artificial conservation for all amino acids present at this position in the MSA. To avoid such a bias, Hopf et al. suggested excluding positions with a higher gap content (curated MSA). Even though this process enhances the performance of a model, it can drastically reduce the number of accessible positions for prediction.

As depicted in Sec. 2.1, the influence of the encoding on the performance of a model turned out to almost vanish for high-*N* data situations. In contrast, we experienced a strong performance-encoding dependency for low-*N* engineering tasks. Here, models based on encoding techniques that provide inherent knowledge of the protein, and can thus counteract the loss of information to a certain extent, were superior to models trained on more general encoded sequences. As maximizing the number of positions for predicting, i.e., including positions with a high gap content when inferring the probabilistic model, and maximizing the quality of the encoding, i.e., excluding positions with a high gap content when inferring the probabilistic model, are counteracting, it is necessary to balance both processes with respect to the underlying data situation. Thus, increasing the size of the training dataset allows to include positions with higher gap content when inferring the probabilistic model, as the information is no longer predominantly sourced from the encoding but from the experimental data.

To quantify the quality of an encoding and thus investigate the influence of the curation of the MSA on the model performance, two statistical models were generated and compared. The parameters of one model were derived based on the “raw MSA”, while the other model is based on a curated MSA according to Hopf et al. [31]. The value of *ρ*_raw_ was determined by ranking all variants that were excluded by the curated model due to the high gap content. The performance of the curated model represents the performance obtained after randomly sampling the same number of variants as excluded variants with effective positions and ranking them. The sampling process was repeated ten times to calculate the mean performance and the standard error on the mean. Fig. 7 highlights the normalized absolute performance difference of both models. In general, using a curated MSA for parameter inference led to a model with enhanced performance. The datasets BLAT ECOLX Palzkill 2012 and HG FLU Bloom2016 marked in orange are an exception. A closer look showed that many positions had a gap content of slightly more than 30 %, but these were not considered for modeling and led to a decreased performance of the statistical model. In such cases, softening this hard threshold and setting it to, e.g., 40 % is superior. If a threshold of 40 % is used, it still holds that the process of curation increases the model performance for both datasets compared to a raw MSA.

**Fig. 7:**
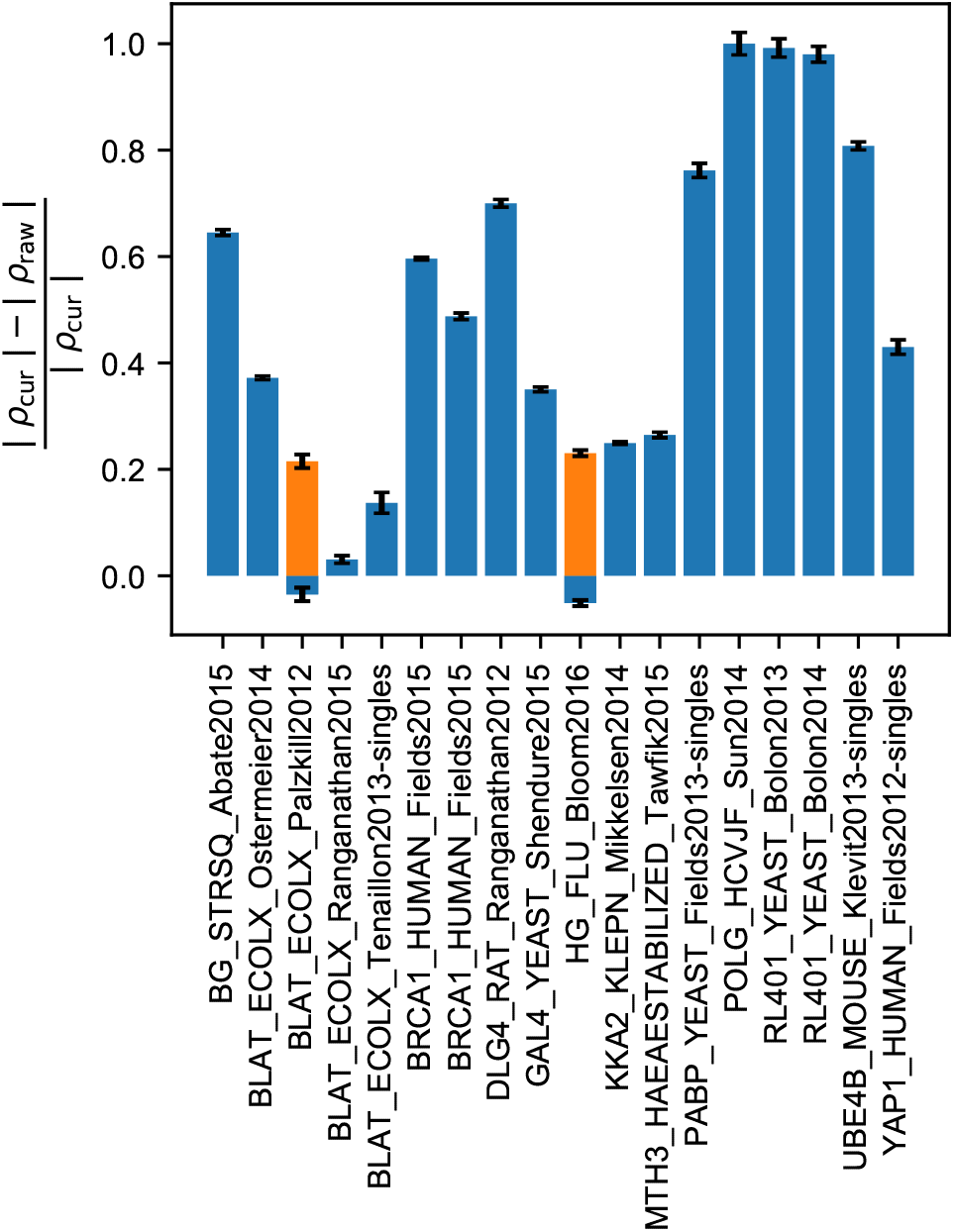
Statistical model performances on curated and non-curated multiple sequence alignments (MSAs). Difference in absolute Spearman’s coefficient of correlation of the statistical models constructed on a curated |*ρ*_cur_| (blue: maximum gap content of 30 % and orange: maximum gap content of 40 %) and non-curated |*ρ*_raw_| (maximum gap content of 100 %) MSA, respectively, achieved when ranking variants according to their fitness label. While the performance of the raw model was estimated on ranking the variants excluded by the curated model, *ρ*_cur_ was obtained as average performance on ranking the same number of variants. The indicated error represents the standard error of the mean.

## 3 Discussion and Conclusion

Protein engineering using directed evolution or (semi-)rational approaches usually results in several tens to hundreds of protein variants that can be genotyped by sequencing to link their measured fitness to an associated sequence. Since establishing a high-throughput screening system or generating a fully diverse protein library requires enormous time and money, data-driven methods are increasingly being integrated into protein engineering campaigns to accelerate the optimization process and navigate through the protein sequence space more efficiently. However, these methods frequently require a large amount of sequence-fitness data, which is often not accessible in practice.

Motivated by this, we developed a hybrid model and compared systematically its performance for different engineering tasks, namely predicting a variant fitness rank from sequence after training (i) on varying dataset sizes ranging from sparse to rich (80 % of the dataset) and (ii) on a varying size of a training dataset consisting of single substituted variants and predicting higher substituted variants. We found our hybrid model to be superior in ranking variants correctly according to their associated fitness labels for (i) and (ii) in regions 50 ≤*N*_train_ ≤250 compared to all other state-of-the-art methods investigated (EVmutation [31], ECNet [24], eUniRep [23], and UniRep [22]) and thus to be especially suited to support protein engineering campaigns even at sparse data situations. In addition, similar to the other supervised methods tested, the hybrid model shows a gain in performance when enlarging the dataset provided for training and can reach similar performance when training on 80 % and testing the remaining 20 % compared to the other high-performance methods tested, i.e., ECNet and eUniRep. This does not only apply when ranking single-substituted variants, but does also hold when predicting the fitness of higher substituted variants after training on single-substituted variant data. With our hybrid model we provide one approach for protein engineers that achieves high performances in predicting the fitness of a protein sequence for both sparse and rich data situations, as well as for predicting the fitness of single and higher-substituted variants. Moreover, within our method we apply a technique to generate low-dimensional encoded sequences that requires less computational effort to train the regressor compared to the other methods presented, which provide higher dimensional encoded sequences to the regressor.

Sequence representations for model training were generated by integrating parameters of a statistical model inferred based on an MSA in the encoding process. The hybrid model has this statistical model as its backbone, which does not require any sequence-fitness data to rank variants. Moreover, it can be adapted to the system by supervised training, which increases the ranking performance. Here, it turned out that the predictions of the supervised term gained weight as the size of the training dataset grew. The underlying encoding corresponds to an “amino acid to scalar” mapping, such that in general significantly fewer parameters need to be adjusted during supervised training compared to the other encoding techniques examined, which use either an “amino acid to vector” or “sequence to high-dimensional vector” mapping. This encoding technique shortens the computational effort and also reduces the memory requirements, while maintaining comparable performances to the state-of-the-art methods. Instead of generating the encoded sequence using a complex recurrent neural network, the “amino acid to scalar mapping” represents a much more transparent and comprehensible encoding scheme.

Furthermore, we found our hybrid model nearly independent of the regularized linear regression method used, unlike methods using higher-dimensional sequence representations [22, 23]. Especially models based on one-hot encoding showed a tendency to overfit using *L*^2^-regularized linear regression (Supplementary Tabs. 3 - 7). Besides, for the datasets studied, we found the general trend that Ridge regression is less regularizing than *L*^1^ regularizing Lasso regression.

As already shown in previous studies [40–42], the derived statistical parameters can be linked to the structure, for example, and thus be interpreted descriptively. This facilitates to draw conclusions about underlying physical principles, which can be used in the further course of the optimization process. It turned out that our approach can be used not only for interpolation tasks (predicting the fitness of a substitution at a position known in the training dataset), but is also valid for positional extrapolation (predicting substitutions at unknown positions). Furthermore, it masters (re-)combination, i.e., substitutional extrapolation tasks, at low data density, because the encoding includes the explicit interactions of two residues, so that nonlinear interactions, which were frequently described in literature [43–46], can also be captured. Finally, it could be shown that curating an MSA always has to be reconciled with the intention of the model (best performance versus providing as many effective positions as possible) and the data situation.

In summary, we found that unlike high-dimensional encoding techniques combined with deep or multiplicative recurrent neural networks, hybrid evolutionary probability and supervised regression modeling using machine learning based on low-dimensional sequence representations not only provides very efficient training, but also usually benefits from better generalization behavior than more complex methods when trained on datasets with a small number of variants. These results demonstrate the suitability of our hybrid model for realistic protein engineering tasks with tens to hundreds of variants, as regularly encountered in academia and industry.

## 4 Methods

### 4.1 Additive Model

Let *I*_***σ***_ be the set containing all position indices *i* of the variant ***σ***, where a substitution has been introduced. Then, the additive model *a* can be defined as

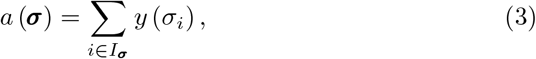

with *y* (*σ*_*i*_) being the measured fitness of the single substituted variant possessing the amino acid *σ*_*i*_ at *i*. A good performance of the additive model suggests that there are no dominant epistatic effects in the recombinants.

### 4.2 Construction of Multiple Sequence Alignments

Unless otherwise noted, the MSAs were generated as follows. First, we performed a jackhmmer search [47] (HMMER version 3.3.2) for extracting sequences from the UniRef100 database (release 2021 03, June 09, 2021) [48] using a bit score of 0.5*L*, i.e., half of the target sequence length for all datasets except for avGFP Kondrashov2016 and BRCA1 HUMAN Fields2015, where a bit score of 0.4 was used to increase the size of the MSA. Next, we post-processed the MSAs, similar to Hopf et al. [31], by excluding (i) all gap positions of the wild-type, (ii) positions with more than 30 % gaps, and (iii) sequences with more than 50 % gaps. The resulting MSA sizes for each dataset are listed in the Supplementary Information (Supplementary Tab. 1).

### 4.3 EVmutation

First, an MSA was generated as described in Sec. 4.2. Based on this MSA, the software plmc (release May 16, 2018), written by John Ingraham in Debora Marks’ lab at the Harvard Medical School, which is available at https://github.com/debbiemarkslab/plmc, was used to generate the matrices ***h***_*i*_ and ***J*** _*ij*_. For inference, the regularization parameters le and lh were set to 0.2 *·* (*N*_sites_ − 1) and 0.01, respectively [31]. *N*_sites_ represents the number of effective sites in the model. The iterative process was run until a minimization success was achieved or 3500 iteration steps were reached. As performance metric, Spearman’s rank correlation coefficient was calculated between the true fitness value and the difference in statistical energy Δ*E* [31] of the variant and the wild-type

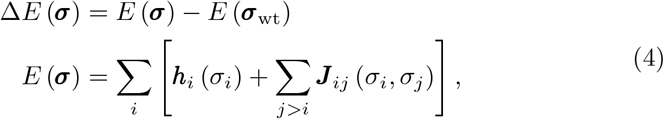

with ***σ*** = (*σ*_1_, *σ*_2_, …, *σ*_*L*_) and *σ*_*i*_ being a representative of one of the 20 proteinogenic amino acids, ***h***_*i*_ (*σ*_*i*_) being an estimate of the local field on position *i*, when occupied with *σ*_*i*_, ***J*** _*ij*_ (*σ*_*i*_, *σ*_*j*_) being an estimate of the coupling between the positions *i* and *j*, when occupied with *σ*_*i*_ and *σ*_*j*_, respectively, and *i, j ∈* [1, *N*_sites_].

### 4.4 Models Using Regularized Linear Regression

Since the focus of this study was to compare different encoding techniques, the supervised models were constructed using an *L*^2^-regularized linear model, whose parameters were set by applying fivefold cross-validation on the training data. For adjusting the regularization parameter, a grid consisting of 100 values sampled from [10^−6^, 10^6^] was searched for the value that minimizes the loss function. To determine the performance of a model in terms of *ρ*, ten different training/test splits were generated for each model. In the following, the encoding strategies used are presented.

#### 4.4.1 One-hot

In one-hot encoding, the canonical amino acids are represented by a set of binary vectors that are linearly independent. As a result, the encoded sequence corresponds to a series of twenty-dimensional binary vectors. As proved by Hsu et al., the coefficients of a position must sum to zero when training an *L*^2^-regularized linear regressor with one-hot encoded sequences [26]. Therefore, there are only *q* − 1 independent coefficients left at each position *c*_*i*_.

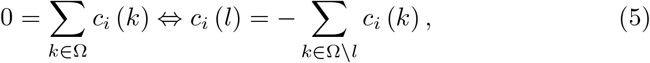

with Ω being the set of all canonical amino acids. Interestingly, in all *L*^2^-regularized one-hot models generated, *l* was found to be the amino acid of the wild-type at position *i*. Therefore, the fitness prediction of an unknown variant with one substitution at position *i* corresponds to the impact of the wild-type amino acid absence, representing the (negative) sum of all amino acid exchange coefficients given in the training dataset at this position.

#### 4.4.2 UniRep

Instead of piecewise assigning each amino acid of a sequence to a scalar or a vector of scalars, respectively, Alley et al. presented in 2019 an approach that takes the whole sequence as input and converts it to a 1900-dimensional vector by passing it through a recurrent neural network [22, 23]. This encoding is based on a hybrid architecture called multiplicative long-short term memory (mLSTM) for conditional sequence modeling tasks that combines LSTM cells for gating the information flow and multiplicative RNN architectures for enabling the use of flexible input-dependent transition functions [49]. This model allows significant improvement in extracting sensitive sentiment representations from text documents and has been used to learn statistical representations of proteins, called UniRep [22] from more than 24 million sequences given in the UniRef50 database [50]. The parameters of the mLSTM network for creating these 1900-dimensional UniRep vectors have been published [22, 51]. In this work, we generated UniRep encoded sequences using the implementation of the *in silico* protein engineering pipeline presented by Favor et al. [51].

#### 4.4.3 eUniRep

Biswas et al. have shown that fine-tuning the recurrent neural network used for generating UniRep encoded sequences with – to the case of study – evolutionary related proteins results in representations, called evotuned UniRep or simply eUniRep, that are superior compared to UniRep [23]. For evotuning, the same procedure as described by Biswas et al. was applied [23]. In contrast, the MSA was obtained by performing a jackhmmer (version 3.3.2) [47] search on the UniRef50 database (release 2021 03, June 09, 2021) [48].

#### 4.4.4 Hybrid Model

At first, the matrices ***h***_*i*_ and ***J*** _*ij*_ were generated as described in Sec. 4.3. For constructing a hybrid model, the linear combination *g* with coefficients *α*_*k*_ ∈ ℝ and *f* (*σ*_0_) = 1 is introduced.

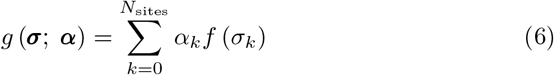

***α*** is obtained by *L*^2^-regularized linear regression. Further, the encoding function *f* is defined as:

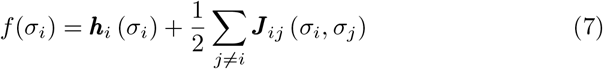

The hybrid model *m*_hybrid_ can be understood as blend of the difference in total energy between ***σ*** and the wild-type sequence ***σ***_wt_ as defined by Hopf et al. [31] with the output of a linear model trained on evolutionary probability model-based encoded sequences. The contribution of the terms is controlled by the parameters *β*_1_, *β*_2_ *∈* [0, 1].

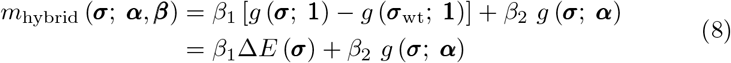

While *β*_1_ = 0 and *β*_2_ ≠ 0 only incorporates the linear model, *β*_1_ ≠ 0 and *β*_2_ = 0 corresponds to the EVmutation approach [31]. *β*_1_ and *β*_2_ are adjusted such that *m*_hybrid_ maximizes the Spearman’s absolute rank correlation coefficient between the true and predicted labels of the training sequences applying five-fold cross-validation. The workflow of the hybrid model algorithm implemented in computer code is shown in the Supplementary Information (Supplementary Fig. 1).

#### 4.4.5 Pure ML Model

The pure ML model corresponds to a hybrid model with *β*_1_ = 0 and *β*_2_ = 1, i.e., to a pure supervised regression-based model.

### 4.5 ECNet

ECNet [24] is a deep learning framework for protein engineering that integrates global and local sequence representations, i.e., amino acid sequence encoding. The global encoding is generated from a language transformer model as formerly implemented in TAPE [52] and used for capturing semantic-rich sequence representations from the protein sequence database Pfam [53]. The pre-trained weights of the TAPE model are used to encode a 768-dimensional vector representation for each amino acid at each sequence position, resulting in a variant sequence being encoded by a matrix of dimension *L* × 768. The local sequence representation is generated by a DCA model similar to the encoding presented herein and is derived from the evolutionary context provided by database-driven MSA construction. However, instead of each amino acid of a sequence being represented by the sums of the local and coupling terms relative to the wild-type couplings at that position, each amino acid is represented by the full *L* + 1 long vector representation including the local and all position-related coupling terms. Thus, a variant sequence encoding results in a matrix of dimension *L* × (*L*+1). Further, dimensions of the amino acid vectors of length (*L* + 1) are reduced by dimensionality reduction to fixed vector lengths *d*, with *d* < *L*. Both sequence representations are fed into a bidirectional short-term memory (BiLSTM) network (a two-layer, fully connected neural network) with the corresponding fitness labels for supervised learning and inference.

We set up files required for running ECNet as described in the respective GitHub repository available at https://github.com/luoyunan/ECNet and generated local features (coupling files) using HHblits [54] for sequence alignment and CCMPred [55] for computing the coupling terms. For constructing alignments with HHblits provided in the HH-suite3, we used Uniclust30 [48] (release June 2020) as sequence database setting parameters for searching homologous sequences as described by Luo et al. [24] (three search iterations, maximum pairwise sequence identity set to 99 %, minimum coverage with query set to 50 %). Subsequently, we inferred couplings from the resulting MSAs using CCMPred and provided the file as local features for supervised training of ECNet. Instead of running ECNet in ensemble mode (by default, three models are created independently and the prediction is averaged), we limited the predictions to a single model and adapted the provided ECNet framework to accept defined data samples for training (randomly sampled learning and validation sets with sizes of 0.8 · *N*_train_ and 0.2 · *N*_train_, respectively) and testing (*N*_total_ − *N*_train_).

## Supporting information

Manuscript SI

## Acknowledgements

This work was partially funded from the Bundesministerium für Bildung und Forschung (BMBF) project (FKZ: 01DJ20014). Model performance studies were performed with computing resources granted by JARA-HPC from RWTH Aachen University under project p0020054. We thank André Breuer for the lively discussions. We thank Dr. David Medina-Ortiz and Prof. Christoph Bannwarth for reading the manuscript.

## Author Contributions

M.D.D. and U.S. supervised the project and acquired research funding. A.-M.I. and N.E.S. performed all experiments, analyzed the results, wrote computer code, and drafted the manuscript. All authors contributed to the revision of the manuscript.

## Data Availability

All SSM datasets and the PABP DMS dataset were taken from the supplementary information provided by Hopf et al. [31]. The avGFP DMS dataset was taken from the additional information provided by Sarkisyan et al. [56]. All protein datasets (SSM and DMS datasets) were further curated to include only positional substitutions considered for DCA-based modeling (30 % maximum gap content in the MSA) as presented in the Supplementary Information (Supplementary Tab. 1). Datasets generated and/or analyzed during the current study are also available from the corresponding author upon reasonable request.

## Computer Code

Encoded datasets, computer code (written in Python 3), and commands for MSA construction, DCA-based encoding, and hybrid modeling are publicly available on GitHub at https://github.com/Protein-Engineering-Framework/HybridModel.

