## Supplementary material for "A hybrid model combining evolutionary probability and machine learning leverages data-driven protein engineering": Manuscript SI

### Supplementary Figures

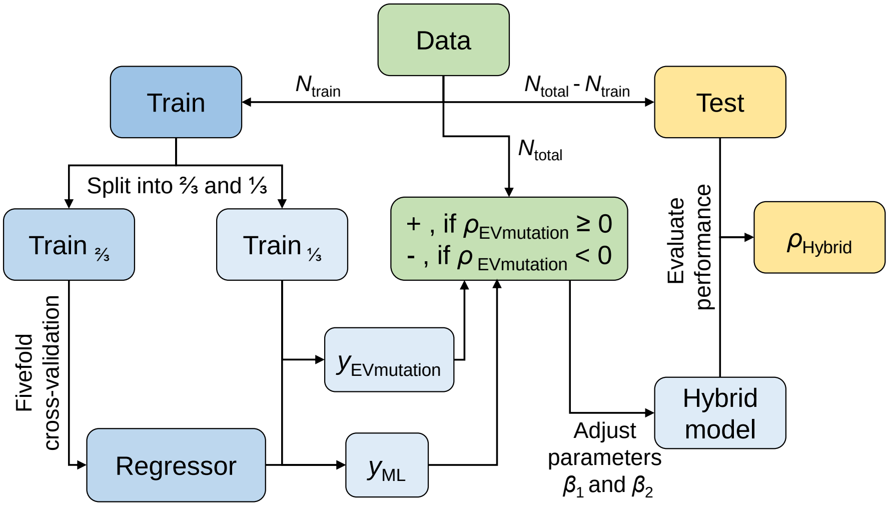


**Supplementary Figure 1**: **Workflow for constructing a hybrid model for prediction of protein fitness from sequence.** The full dataset (green) is split into sets for supervised regression model training (blue) and testing (yellow). The linear regression model is trained on 2/3 of the training data using fivefold cross-validation (CV). The remaining 1/3 of the training data are subsequently used to adjust the contribution of the supervised (ML) and the statistical model (EVmutation) to the hybrid model. The model contributions are adjusted by finding the set $\{\beta_1, \beta_2\}$ that maximizes Spearman's $\rho$ on $\text{Train}_{1/3}$. Finally, the performance of the hybrid model's predictions is evaluated as the Spearman correlation between the observed fitness values and the predicted fitness values of the test set ($\rho_{\text{Hybrid}}$).

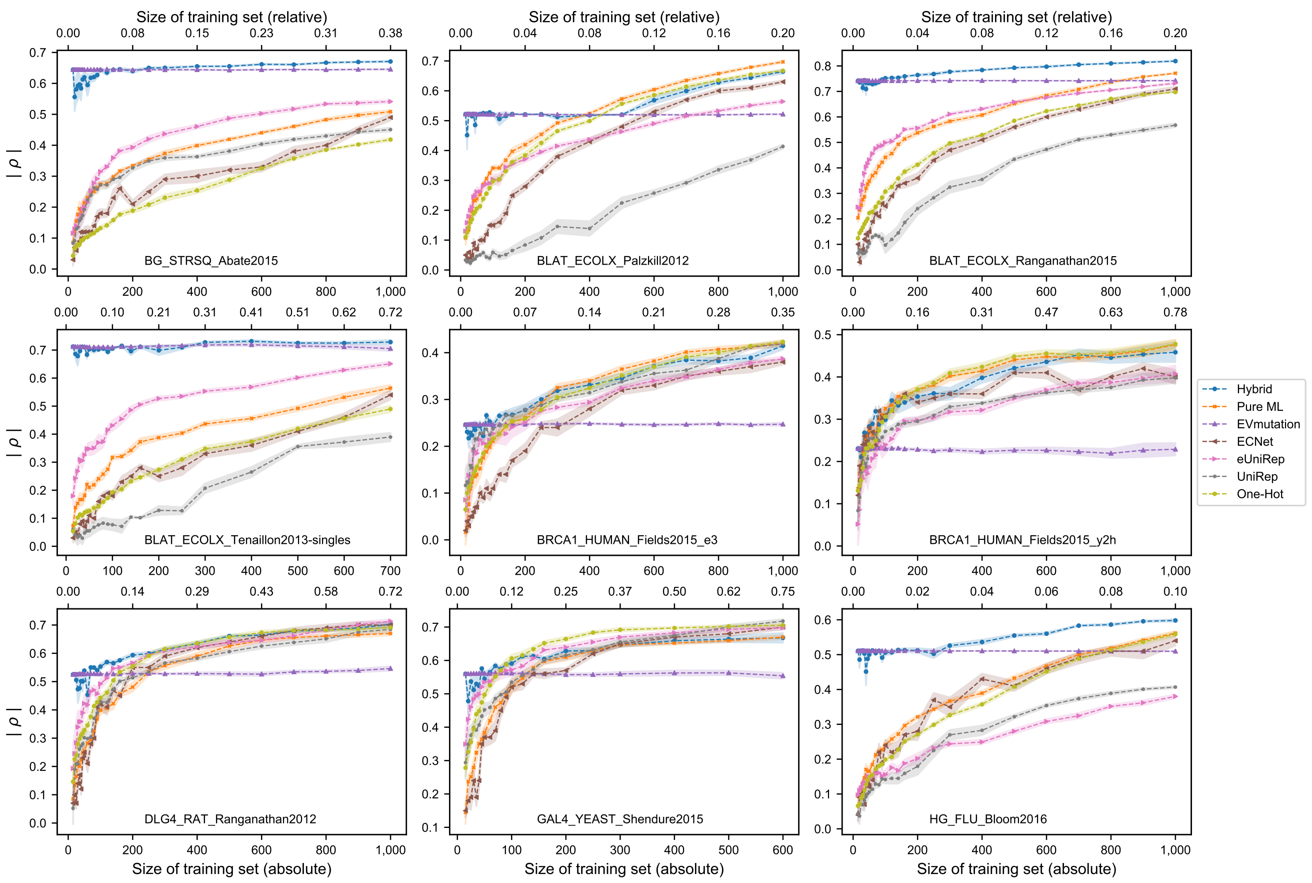


**Supplementary Figure 2**: **Performance of different methods for individual datasets for ranking the variants according to their fitness label at different training dataset sizes.** The rank coefficients represent the averaged performance of ten different splits when ranking the holdout variants according to their fitness label. Colored areas indicate the standard error of the mean. The test performances of the hybrid, pure ML, EVmutation [1], ECNet [2], eUniRep [3], UniRep [4], and one-hot model are compared.

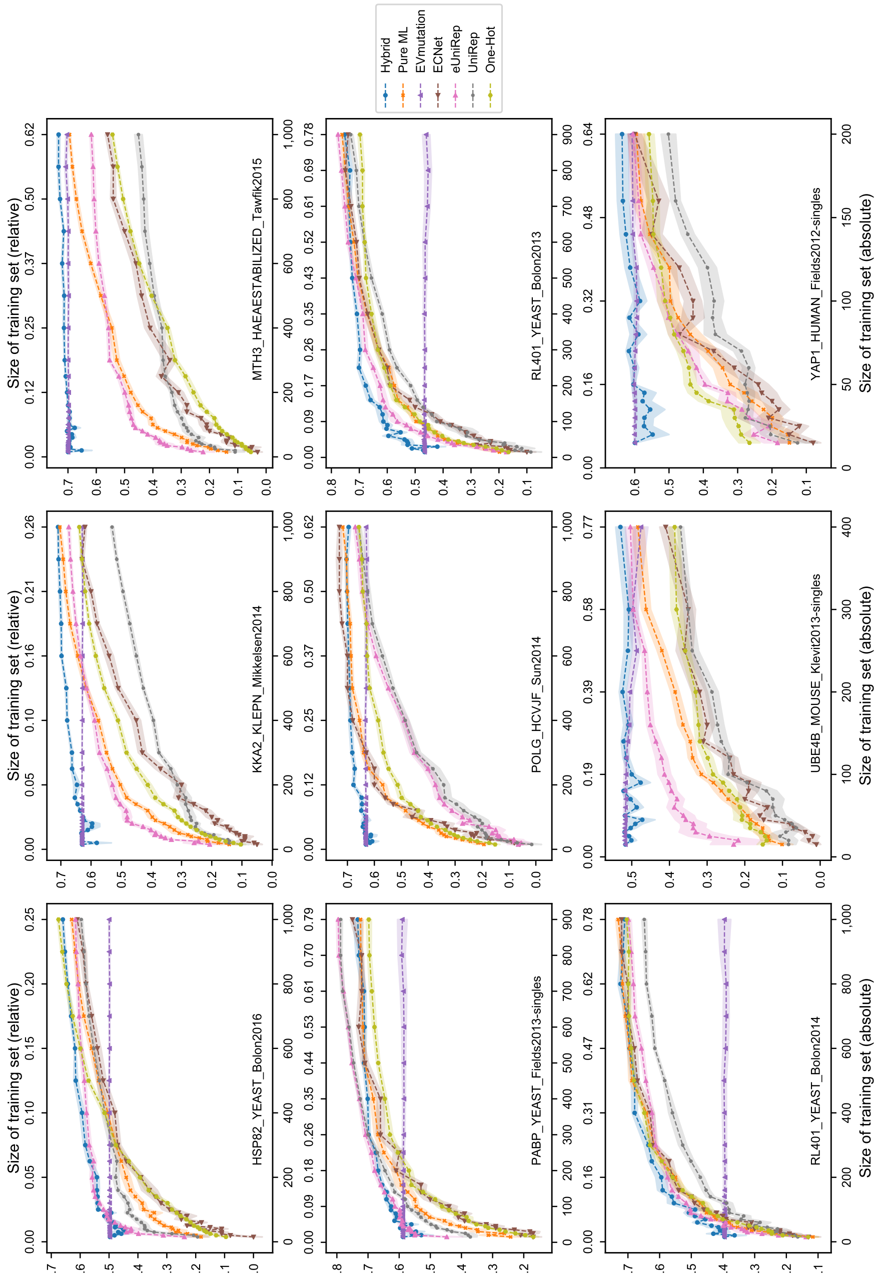


**Supplementary Figure 2** (cont.)

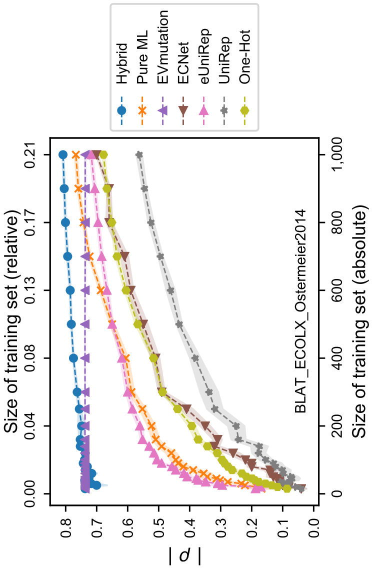


**Supplementary Figure 2** (cont.)

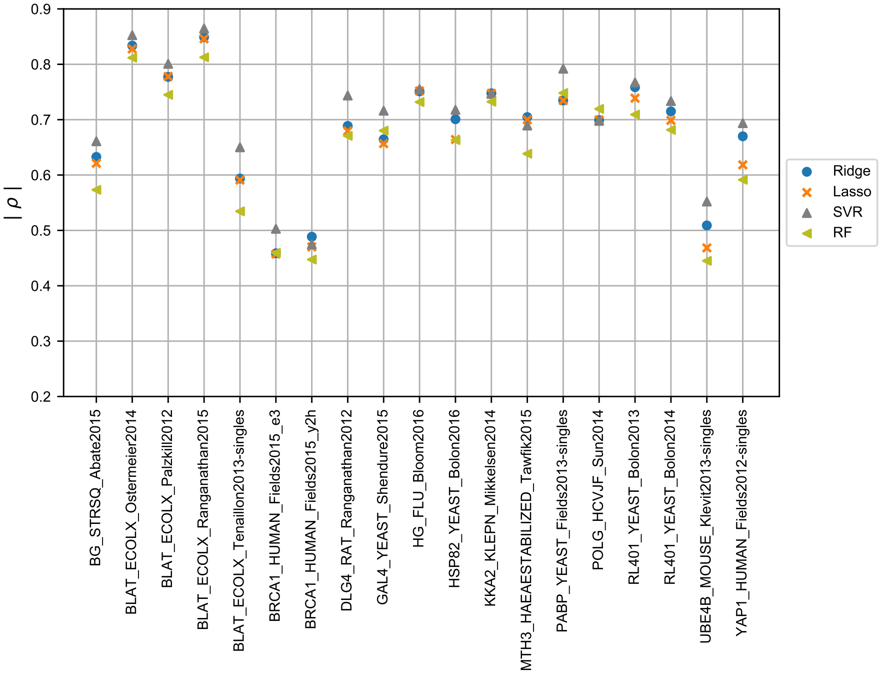


**Supplementary Figure 3**: **Performance of different regression methods for ranking the variants according to their fitness label for all SSM datasets.** The absolute Spearman rank correlation coefficients observed on 19 SSM datasets for different pure ML-based models when trained and tested on 80 % and 20 % of the data, respectively. The performances of the linear regression models (Ridge, Lasso, and ordinary least squares (OLS) regression) as well as the nonlinear models (support vector regression (SVR) and random forest regression (RF)) represent the mean of 10 different splits. For adjusting the hyperparameters of the models, the following parameter ranges were used: Ridge/Lasso: $\alpha$ (100 values with logarithmically spaced distance ranging from $10^{-6}$ to $10^{6}$), SVR: $C$ (50 values with logarithmically spaced distance ranging from $10^{-3}$ to $10^{3}$) and the tube penalty parameter $\epsilon$ (50 values with logarithmically spaced distance ranging from $10^{-4}$ to $10^{2}$), and RF: number of estimators ($\{1, 5, 10, 20, 50, 100, 200, 500, 1000\}$) and the maximum number of used features ($\{$all features, $\sqrt{\text{all features}}$, $\log_2$ all features$\}$).

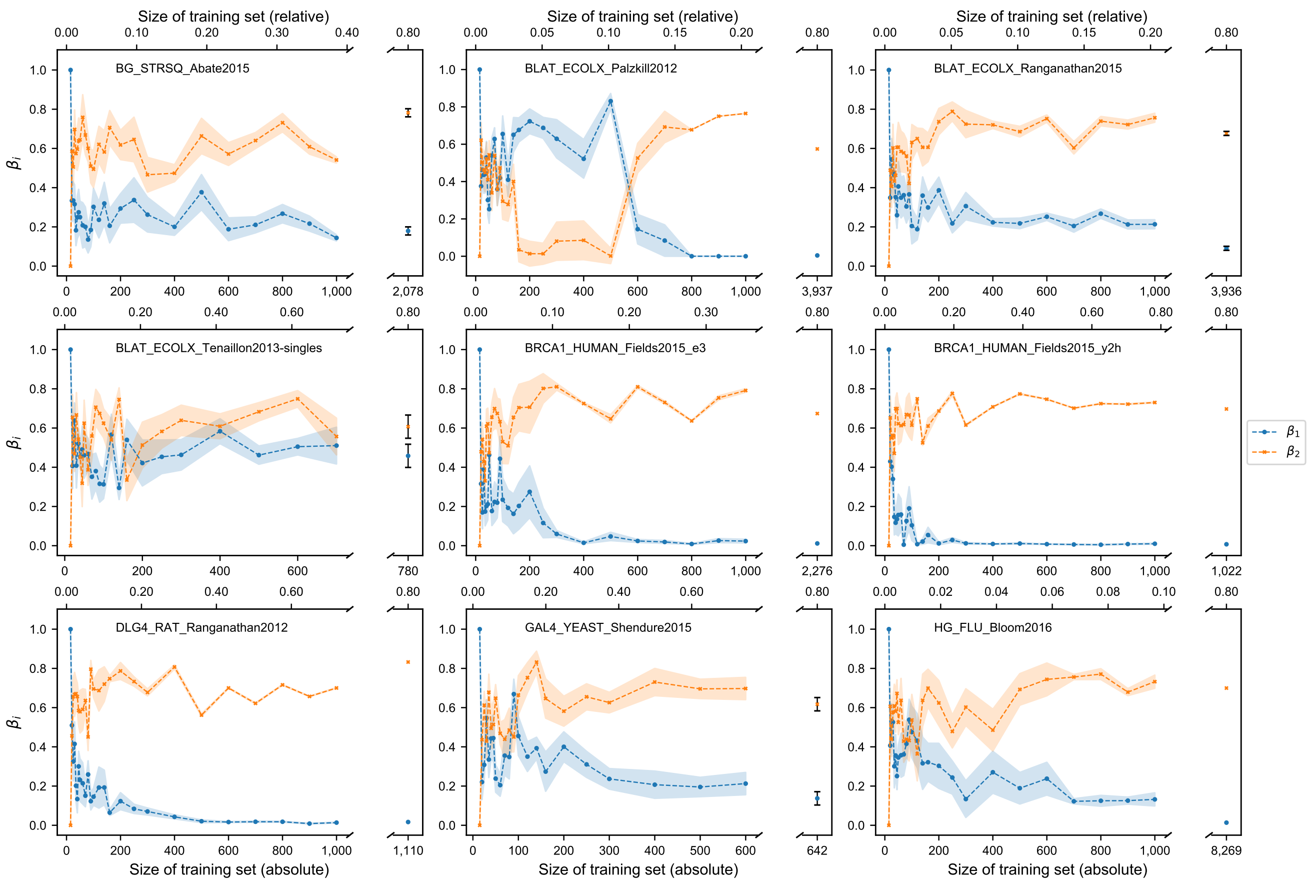


**Supplementary Figure 4**: **Influence of the DCA and ML model on the predicted fitness of the hybrid model at different training dataset sizes of 19 SSM datasets.** The hybrid model results after combining both models with their individual weights, $\beta_1$ (DCA model contribution) and $\beta_2$ (ML model contribution), and is constructed only if at least 20 % of the full dataset remains for prediction. While the dots represent the mean of ten different splits, the colored areas represent the standard error of the mean.

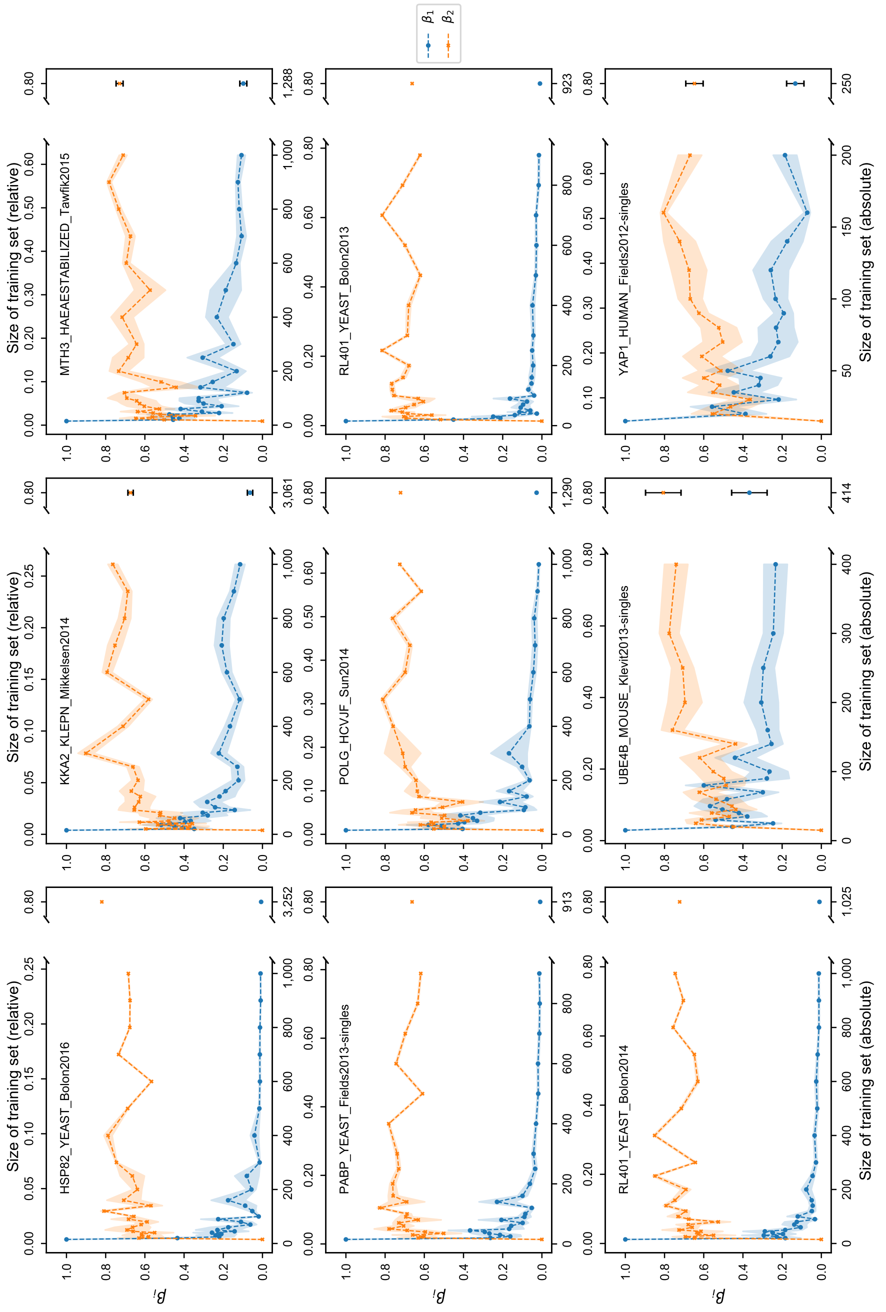


**Supplementary Figure 4** (cont.)

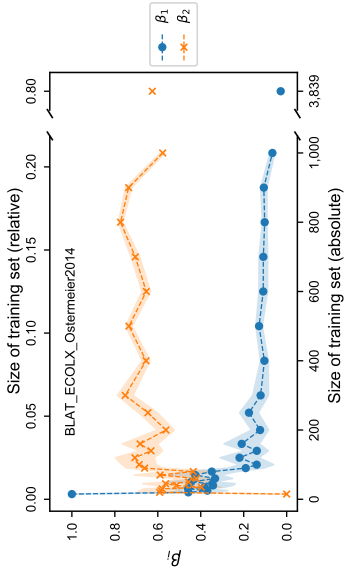


**Supplementary Figure 4** (cont.)

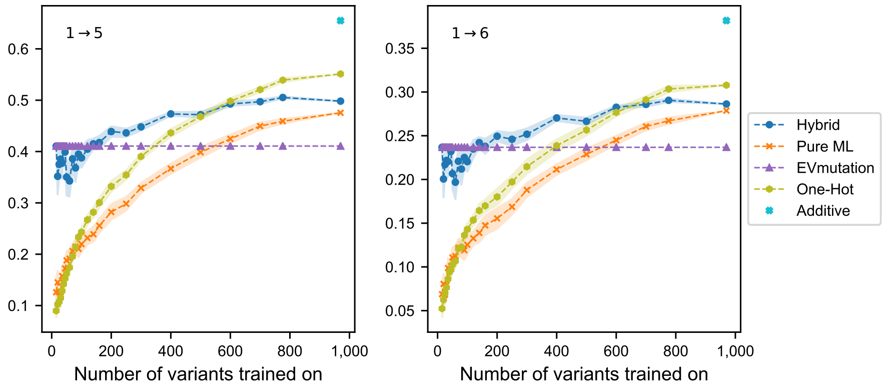


**Supplementary Figure 5**: **Performance of different methods for ranking recombinants of the green fluorescent protein according to their fitness label.** Spearman's rank coefficient of correlation $\rho$ was determined after ranking multiple substituted variants with five ($1 \rightarrow 5$) and six ($1 \rightarrow 6$) simultaneous substitutions of the green fluorescent protein from *Aequorea victoria* as a function of the number of single substituted variants in the training dataset. The colored area represents the standard error of the mean.

### Supplementary Tables

**Supplementary Table 1: Overview of datasets used to derive model parameters through constructed multiple sequence alignments (MSA), MSA post-processing, and direct coupling analysis (DCA).** 'MSA' defines the number of sequences in each MSA, '$N_{\text{sites}}/L$' defines the ratio of effective sites remaining for DCA and target sequence length at maximum gap contents of 30 %, and '$N$' defines the number of remaining variant-fitness pairs used for modeling that corresponds to the effective sites of the MSA after post-processing. All single-saturation mutagenesis (SSM) and the PABP_YEAST_Fields2013 deep mutational scanning dataset were taken from Supplementary Table 2 of Hopf et al. [1]; the avGFP deep mutational scanning dataset was taken from Sarkisyan et al. [5].

| Dataset | MSA | $N_{\text{sites}}/L$ | $N$ |
|---|---|---|---|
| avGFP_Kondrashov2016 [5] | 1015 | 0.90 | 32610 |
| BG_STRSQ_Abate2015 [6] | 109098 | 0.91 | 2598 |
| BLAT_ECOLX_Ostermeier2014 [7] | 27258 | 0.92 | 4799 |
| BLAT_ECOLX_Palzkill2012 [8] | 27258 | 0.92 | 4922 |
| BLAT_ECOLX_Ranganathan2015 [9] | 27258 | 0.92 | 4921 |
| BLAT_ECOLX_Tenaillon2013-singles [10] | 27258 | 0.92 | 975 |
| BRCA1_HUMAN_Fields2015_e3 [11] | 26529 | 0.57 | 2846 |
| BRCA1_HUMAN_Fields2015_y2h [11] | 26529 | 0.57 | 1278 |
| DLG4_RAT_Ranganathan2012 [12] | 746067 | 0.88 | 1388 |
| GAL4_YEAST_Shendure2015 [13] | 144154 | 0.72 | 803 |
| HG_FLU_Bloom2016 [14] | 63771 | 0.96 | 10337 |
| HSP82_YEAST_Bolon2016 [15] | 51619 | 0.94 | 4065 |
| KKA2_KLEPN_Mikkelsen2014 [16] | 5581 | 0.77 | 3827 |
| MTH3_HAEAESTABILIZED_Tawfik2015 [17] | 66125 | 0.91 | 1611 |
| PABP_YEAST_Fields2013-singles [18] | 770434 | 0.95 | 1142 |
| PABP_YEAST_Fields2013-singles-doubles [18] | 770434 | 0.95 | 34913 |
| POLG_HCVJF_Sun2014 [19] | 14351 | 0.99 | 1613 |
| RL401_YEAST_Bolon2013 [20] | 78384 | 0.93 | 1154 |
| RL401_YEAST_Bolon2014 [21] | 78384 | 0.93 | 1282 |
| UBE4B_MOUSE_Klevit2013-singles [22] | 78018 | 0.67 | 518 |
| YAP1_HUMAN_Fields2012-singles [23] | 193046 | 0.88 | 313 |

**Supplementary Table 2: Comparison of different regression methods when ranking the variants according to their fitness label for all SSM datasets.** Mean, minimum, median, and maximum absolute Spearman rank correlation coefficients observed on 19 different datasets for the regressors (pure ML models) under investigation when trained and tested on 80 % and 20 % of the data, respectively. The errors on the median, minimum, as well as maximum value correspond to the standard error of the mean obtained from 10 different splits for the associated dataset.

| **Regressor** | **Mean** | **Median** | **Minimum** | **Maximum** |
|---|---|---|---|---|
| Ridge | 0.68 | $0.70 \pm 0.01$ | $0.46 \pm 0.01$ | $0.85 \pm 0.01$ |
| Lasso | 0.67 | $0.70 \pm 0.01$ | $0.46 \pm 0.01$ | $0.85 \pm 0.01$ |
| OLS | 0.68 | $0.70 \pm 0.01$ | $0.45 \pm 0.01$ | $0.86 \pm 0.01$ |
| SVR | 0.71 | $0.72 \pm 0.01$ | $0.48 \pm 0.03$ | $0.87 \pm 0.01$ |
| RF | 0.65 | $0.68 \pm 0.01$ | $0.45 \pm 0.03$ | $0.81 \pm 0.01$ |

**Supplementary Table 3: One-Hot model performance on the SSM datasets.** Achieved Spearman's $\rho$ of observed and predicted fitness values on the individual SSM datasets for training (80 %) and testing (20 %) using Lasso and Ridge regression. The error given corresponds to the standard error of the mean obtained from 10 different splits for the associated dataset.

| Dataset | Lasso | | Ridge | |
|---|---|---|---|---|
| | $\rho_{\text{train}}$ | $\rho_{\text{test}}$ | $\rho_{\text{train}}$ | $\rho_{\text{test}}$ |
| BG_STRSQ_Abate2015 | $0.74 \pm 0.02$ | $0.51 \pm 0.04$ | $0.96 \pm 0.02$ | $0.54 \pm 0.04$ |
| BLAT_ECOLX_Ostermeier2014 | $0.86 \pm 0.01$ | $0.76 \pm 0.02$ | $1.00 \pm 0.01$ | $0.75 \pm 0.02$ |
| BLAT_ECOLX_Palzkill2012 | $0.83 \pm 0.02$ | $0.75 \pm 0.01$ | $1.00 \pm 0.00$ | $0.74 \pm 0.01$ |
| BLAT_ECOLX_Ranganathan2015 | $0.84 \pm 0.02$ | $0.77 \pm 0.02$ | $1.00 \pm 0.00$ | $0.76 \pm 0.02$ |
| BLAT_ECOLX_Tenaillon2013-singles | $0.77 \pm 0.03$ | $0.42 \pm 0.06$ | $0.98 \pm 0.01$ | $0.50 \pm 0.04$ |
| BRCA1_HUMAN_Fields2015_e3 | $0.61 \pm 0.04$ | $0.47 \pm 0.03$ | $0.88 \pm 0.02$ | $0.47 \pm 0.03$ |
| BRCA1_HUMAN_Fields2015_y2h | $0.62 \pm 0.02$ | $0.50 \pm 0.08$ | $0.93 \pm 0.03$ | $0.50 \pm 0.08$ |
| DLG4_RAT_Ranganathan2012 | $0.76 \pm 0.02$ | $0.70 \pm 0.06$ | $0.99 \pm 0.02$ | $0.70 \pm 0.06$ |
| GAL4_YEAST_Shendure2015 | $0.82 \pm 0.02$ | $0.71 \pm 0.04$ | $1.00 \pm 0.00$ | $0.71 \pm 0.04$ |
| HG_FLU_Bloom2016 | $0.83 \pm 0.02$ | $0.74 \pm 0.01$ | $1.00 \pm 0.00$ | $0.74 \pm 0.01$ |
| HSP82_YEAST_Bolon2016 | $0.77 \pm 0.01$ | $0.73 \pm 0.02$ | $1.00 \pm 0.00$ | $0.73 \pm 0.02$ |
| KKA2_KLEPN_Mikkelsen2014 | $0.80 \pm 0.02$ | $0.69 \pm 0.03$ | $1.00 \pm 0.01$ | $0.69 \pm 0.03$ |
| MTH3_HAEAESTABILIZED_Tawfik2015 | $0.85 \pm 0.03$ | $0.56 \pm 0.04$ | $0.99 \pm 0.01$ | $0.57 \pm 0.05$ |
| PABP_YEAST_Fields2013-singles | $0.76 \pm 0.01$ | $0.72 \pm 0.04$ | $1.00 \pm 0.01$ | $0.72 \pm 0.04$ |
| POLG_HCVJF_Sun2014 | $0.76 \pm 0.02$ | $0.67 \pm 0.03$ | $0.91 \pm 0.01$ | $0.66 \pm 0.03$ |
| RL401_YEAST_Bolon2013 | $0.79 \pm 0.01$ | $0.71 \pm 0.03$ | $0.96 \pm 0.03$ | $0.70 \pm 0.03$ |
| RL401_YEAST_Bolon2014 | $0.76 \pm 0.01$ | $0.67 \pm 0.04$ | $0.99 \pm 0.02$ | $0.70 \pm 0.03$ |
| UBE4B_MOUSE_Klevit2013-singles | $0.75 \pm 0.25$ | $0.32 \pm 0.14$ | $0.91 \pm 0.04$ | $0.41 \pm 0.09$ |
| YAP1_HUMAN_Fields2012-singles | $0.92 \pm 0.10$ | $0.66 \pm 0.06$ | $0.97 \pm 0.03$ | $0.64 \pm 0.07$ |

**Supplementary Table 4**: **Hybrid model performance on the SSM datasets.** Achieved Spearman's $\rho$ of observed and predicted fitness values on the individual SSM sets for training (80 %) and testing (20 %) using Lasso and Ridge regression. The error given corresponds to the standard error of the mean obtained from 10 different splits for the associated dataset.

| Dataset | Lasso | | Ridge | |
|---|---|---|---|---|
| | $\rho_{\text{train}}$ | $\rho_{\text{test}}$ | $\rho_{\text{train}}$ | $\rho_{\text{test}}$ |
| BG_STRSQ_Abate2015 | $0.71 \pm 0.02$ | $0.67 \pm 0.03$ | $0.73 \pm 0.02$ | $0.68 \pm 0.03$ |
| BLAT_ECOLX_Ostermeier2014 | $0.86 \pm 0.01$ | $0.84 \pm 0.01$ | $0.86 \pm 0.01$ | $0.84 \pm 0.02$ |
| BLAT_ECOLX_Palzkill2012 | $0.80 \pm 0.01$ | $0.77 \pm 0.02$ | $0.80 \pm 0.01$ | $0.77 \pm 0.02$ |
| BLAT_ECOLX_Ranganathan2015 | $0.87 \pm 0.01$ | $0.86 \pm 0.01$ | $0.87 \pm 0.01$ | $0.86 \pm 0.01$ |
| BLAT_ECOLX_Tenaillon2013-singles | $0.75 \pm 0.01$ | $0.71 \pm 0.04$ | $0.78 \pm 0.02$ | $0.72 \pm 0.04$ |
| BRCA1_HUMAN_Fields2015_e3 | $0.52 \pm 0.02$ | $0.44 \pm 0.05$ | $0.54 \pm 0.01$ | $0.45 \pm 0.05$ |
| BRCA1_HUMAN_Fields2015_y2h | $0.49 \pm 0.06$ | $0.43 \pm 0.06$ | $0.54 \pm 0.03$ | $0.47 \pm 0.07$ |
| DLG4_RAT_Ranganathan2012 | $0.70 \pm 0.03$ | $0.68 \pm 0.06$ | $0.70 \pm 0.02$ | $0.69 \pm 0.06$ |
| GAL4_YEAST_Shendure2015 | $0.69 \pm 0.01$ | $0.66 \pm 0.05$ | $0.70 \pm 0.01$ | $0.66 \pm 0.04$ |
| HG_FLU_Bloom2016 | $0.77 \pm 0.01$ | $0.74 \pm 0.01$ | $0.77 \pm 0.01$ | $0.74 \pm 0.01$ |
| HSP82_YEAST_Bolon2016 | $0.70 \pm 0.02$ | $0.68 \pm 0.03$ | $0.72 \pm 0.01$ | $0.71 \pm 0.03$ |
| KKA2_KLEPN_Mikkelsen2014 | $0.77 \pm 0.01$ | $0.75 \pm 0.02$ | $0.77 \pm 0.01$ | $0.75 \pm 0.02$ |
| MTH3_HAEAESTABILIZED_Tawfik2015 | $0.77 \pm 0.01$ | $0.72 \pm 0.03$ | $0.78 \pm 0.01$ | $0.72 \pm 0.03$ |
| PABP_YEAST_Fields2013-singles | $0.76 \pm 0.02$ | $0.73 \pm 0.05$ | $0.76 \pm 0.02$ | $0.73 \pm 0.05$ |
| POLG_HCVJF_Sun2014 | $0.74 \pm 0.01$ | $0.70 \pm 0.03$ | $0.74 \pm 0.02$ | $0.69 \pm 0.03$ |
| RL401_YEAST_Bolon2013 | $0.79 \pm 0.01$ | $0.74 \pm 0.03$ | $0.79 \pm 0.01$ | $0.74 \pm 0.03$ |
| RL401_YEAST_Bolon2014 | $0.74 \pm 0.03$ | $0.70 \pm 0.03$ | $0.76 \pm 0.02$ | $0.71 \pm 0.03$ |
| UBE4B_MOUSE_Klevit2013-singles | $0.59 \pm 0.05$ | $0.49 \pm 0.04$ | $0.63 \pm 0.03$ | $0.51 \pm 0.04$ |
| YAP1_HUMAN_Fields2012-singles | $0.68 \pm 0.03$ | $0.67 \pm 0.07$ | $0.68 \pm 0.03$ | $0.67 \pm 0.07$ |

**Supplementary Table 5**: **Pure ML model performance on the SSM datasets.** Achieved Spearman's $\rho$ of observed and predicted fitness values on the individual SSM sets for training (80 %) and testing (20 %) using Lasso and Ridge regression. The error given corresponds to the standard error of the mean obtained from 10 different splits for the associated dataset.

| Dataset | Lasso | | Ridge | |
|---|---|---|---|---|
| | $\rho_{\text{train}}$ | $\rho_{\text{test}}$ | $\rho_{\text{train}}$ | $\rho_{\text{test}}$ |
| BG_STRSQ_Abate2015 | $0.74 \pm 0.02$ | $0.62 \pm 0.04$ | $0.76 \pm 0.01$ | $0.63 \pm 0.03$ |
| BLAT_ECOLX_Ostermeier2014 | $0.86 \pm 0.01$ | $0.83 \pm 0.02$ | $0.86 \pm 0.01$ | $0.83 \pm 0.02$ |
| BLAT_ECOLX_Palzkill2012 | $0.81 \pm 0.01$ | $0.78 \pm 0.02$ | $0.81 \pm 0.01$ | $0.78 \pm 0.02$ |
| BLAT_ECOLX_Ranganathan2015 | $0.87 \pm 0.01$ | $0.85 \pm 0.01$ | $0.87 \pm 0.01$ | $0.85 \pm 0.01$ |
| BLAT_ECOLX_Tenaillon2013-singles | $0.79 \pm 0.01$ | $0.59 \pm 0.05$ | $0.81 \pm 0.01$ | $0.59 \pm 0.05$ |
| BRCA1_HUMAN_Fields2015_e3 | $0.55 \pm 0.01$ | $0.46 \pm 0.03$ | $0.56 \pm 0.01$ | $0.46 \pm 0.03$ |
| BRCA1_HUMAN_Fields2015_y2h | $0.55 \pm 0.04$ | $0.47 \pm 0.06$ | $0.56 \pm 0.02$ | $0.49 \pm 0.08$ |
| DLG4_RAT_Ranganathan2012 | $0.72 \pm 0.02$ | $0.69 \pm 0.05$ | $0.72 \pm 0.01$ | $0.69 \pm 0.05$ |
| GAL4_YEAST_Shendure2015 | $0.70 \pm 0.02$ | $0.66 \pm 0.04$ | $0.71 \pm 0.01$ | $0.67 \pm 0.04$ |
| HG_FLU_Bloom2016 | $0.78 \pm 0.01$ | $0.75 \pm 0.01$ | $0.78 \pm 0.01$ | $0.75 \pm 0.01$ |
| HSP82_YEAST_Bolon2016 | $0.71 \pm 0.01$ | $0.69 \pm 0.03$ | $0.73 \pm 0.01$ | $0.70 \pm 0.02$ |
| KKA2_KLEPN_Mikkelsen2014 | $0.78 \pm 0.01$ | $0.75 \pm 0.02$ | $0.78 \pm 0.01$ | $0.75 \pm 0.02$ |
| MTH3_HAEAESTABILIZED_Tawfik2015 | $0.81 \pm 0.01$ | $0.70 \pm 0.04$ | $0.82 \pm 0.01$ | $0.71 \pm 0.03$ |
| PABP_YEAST_Fields2013-singles | $0.77 \pm 0.02$ | $0.74 \pm 0.05$ | $0.77 \pm 0.02$ | $0.74 \pm 0.05$ |
| POLG_HCVJF_Sun2014 | $0.75 \pm 0.01$ | $0.70 \pm 0.03$ | $0.75 \pm 0.01$ | $0.70 \pm 0.03$ |
| RL401_YEAST_Bolon2013 | $0.81 \pm 0.01$ | $0.76 \pm 0.04$ | $0.81 \pm 0.01$ | $0.76 \pm 0.04$ |
| RL401_YEAST_Bolon2014 | $0.76 \pm 0.01$ | $0.71 \pm 0.03$ | $0.77 \pm 0.01$ | $0.72 \pm 0.03$ |
| UBE4B_MOUSE_Klevit2013-singles | $0.66 \pm 0.05$ | $0.47 \pm 0.07$ | $0.70 \pm 0.02$ | $0.51 \pm 0.05$ |
| YAP1_HUMAN_Fields2012-singles | $0.68 \pm 0.13$ | $0.62 \pm 0.24$ | $0.73 \pm 0.02$ | $0.67 \pm 0.10$ |

**Supplementary Table 6**: **eUniRep model performance on SSM datasets.** Achieved Spearman's $\rho$ of observed and predicted fitness values on the individual SSM sets for training (80 %) and testing (20 %) using Lasso and Ridge regression. The error given corresponds to the standard error of the mean obtained from 10 different splits for the associated dataset.

| Dataset | Lasso | | Ridge | |
|---|---|---|---|---|
| | $\rho_{\text{train}}$ | $\rho_{\text{test}}$ | $\rho_{\text{train}}$ | $\rho_{\text{test}}$ |
| BG_STRSQ_Abate2015 | $0.70 \pm 0.02$ | $0.58 \pm 0.04$ | $0.75 \pm 0.01$ | $0.61 \pm 0.02$ |
| BLAT_ECOLX_Ostermeier2014 | $0.88 \pm 0.01$ | $0.81 \pm 0.01$ | $0.89 \pm 0.01$ | $0.81 \pm 0.01$ |
| BLAT_ECOLX_Palzkill2012 | $0.82 \pm 0.01$ | $0.70 \pm 0.02$ | $0.83 \pm 0.01$ | $0.71 \pm 0.02$ |
| BLAT_ECOLX_Ranganathan2015 | $0.89 \pm 0.01$ | $0.81 \pm 0.01$ | $0.89 \pm 0.01$ | $0.82 \pm 0.01$ |
| BLAT_ECOLX_Tenaillon2013-singles | $0.77 \pm 0.05$ | $0.58 \pm 0.05$ | $0.83 \pm 0.01$ | $0.65 \pm 0.05$ |
| BRCA1_HUMAN_Fields2015_e3 | $0.55 \pm 0.04$ | $0.42 \pm 0.03$ | $0.62 \pm 0.02$ | $0.45 \pm 0.03$ |
| BRCA1_HUMAN_Fields2015_y2h | $0.52 \pm 0.07$ | $0.42 \pm 0.04$ | $0.62 \pm 0.04$ | $0.46 \pm 0.04$ |
| DLG4_RAT_Ranganathan2012 | $0.77 \pm 0.03$ | $0.67 \pm 0.05$ | $0.83 \pm 0.03$ | $0.72 \pm 0.04$ |
| GAL4_YEAST_Shendure2015 | $0.80 \pm 0.03$ | $0.67 \pm 0.04$ | $0.87 \pm 0.02$ | $0.70 \pm 0.04$ |
| HG_FLU_Bloom2016 | $0.74 \pm 0.01$ | $0.64 \pm 0.02$ | $0.75 \pm 0.01$ | $0.65 \pm 0.02$ |
| HSP82_YEAST_Bolon2016 | $0.72 \pm 0.01$ | $0.66 \pm 0.03$ | $0.74 \pm 0.01$ | $0.67 \pm 0.02$ |
| KKA2_KLEPN_Mikkelsen2014 | $0.82 \pm 0.01$ | $0.72 \pm 0.02$ | $0.84 \pm 0.01$ | $0.73 \pm 0.02$ |
| MTH3_HAEAESTABILIZED_Tawfik2015 | $0.77 \pm 0.02$ | $0.63 \pm 0.04$ | $0.81 \pm 0.02$ | $0.65 \pm 0.04$ |
| PABP_YEAST_Fields2013-singles | $0.89 \pm 0.02$ | $0.78 \pm 0.03$ | $0.91 \pm 0.01$ | $0.80 \pm 0.02$ |
| POLG_HCVJF_Sun2014 | $0.77 \pm 0.02$ | $0.67 \pm 0.04$ | $0.82 \pm 0.01$ | $0.71 \pm 0.04$ |
| RL401_YEAST_Bolon2013 | $0.85 \pm 0.01$ | $0.74 \pm 0.03$ | $0.87 \pm 0.01$ | $0.77 \pm 0.02$ |
| RL401_YEAST_Bolon2014 | $0.79 \pm 0.02$ | $0.68 \pm 0.05$ | $0.82 \pm 0.01$ | $0.71 \pm 0.04$ |
| UBE4B_MOUSE_Klevit2013-singles | $0.67 \pm 0.04$ | $0.44 \pm 0.06$ | $0.77 \pm 0.03$ | $0.52 \pm 0.06$ |
| YAP1_HUMAN_Fields2012-singles | $0.84 \pm 0.07$ | $0.55 \pm 0.11$ | $0.91 \pm 0.03$ | $0.66 \pm 0.09$ |

**Supplementary Table 7**: **UniRep model performance on SSM datasets.** Achieved Spearman's $\rho$ of observed and predicted fitness values on the individual SSM sets for training (80 %) and testing (20 %) using Lasso and Ridge regression. The error given corresponds to the standard error of the mean obtained from 10 different splits for the associated dataset.

| Dataset | Lasso | | Ridge | |
|---|---|---|---|---|
| | $\rho_{\text{train}}$ | $\rho_{\text{test}}$ | $\rho_{\text{train}}$ | $\rho_{\text{test}}$ |
| BG_STRSQ_Abate2015 | $0.59 \pm 0.02$ | $0.47 \pm 0.06$ | $0.66 \pm 0.03$ | $0.52 \pm 0.05$ |
| BLAT_ECOLX_Ostermeier2014 | $0.82 \pm 0.02$ | $0.73 \pm 0.03$ | $0.84 \pm 0.01$ | $0.75 \pm 0.02$ |
| BLAT_ECOLX_Palzkill2012 | $0.75 \pm 0.06$ | $0.61 \pm 0.05$ | $0.78 \pm 0.03$ | $0.64 \pm 0.02$ |
| BLAT_ECOLX_Ranganathan2015 | $0.84 \pm 0.03$ | $0.73 \pm 0.04$ | $0.83 \pm 0.02$ | $0.74 \pm 0.02$ |
| BLAT_ECOLX_Tenaillon2013-singles | $0.61 \pm 0.07$ | $0.37 \pm 0.06$ | $0.66 \pm 0.03$ | $0.42 \pm 0.06$ |
| BRCA1_HUMAN_Fields2015_e3 | $0.56 \pm 0.02$ | $0.46 \pm 0.04$ | $0.62 \pm 0.01$ | $0.48 \pm 0.04$ |
| BRCA1_HUMAN_Fields2015_y2h | $0.53 \pm 0.03$ | $0.44 \pm 0.05$ | $0.57 \pm 0.02$ | $0.43 \pm 0.06$ |
| DLG4_RAT_Ranganathan2012 | $0.75 \pm 0.04$ | $0.66 \pm 0.06$ | $0.81 \pm 0.02$ | $0.70 \pm 0.05$ |
| GAL4_YEAST_Shendure2015 | $0.87 \pm 0.03$ | $0.71 \pm 0.05$ | $0.88 \pm 0.02$ | $0.73 \pm 0.03$ |
| HG_FLU_Bloom2016 | $0.72 \pm 0.02$ | $0.63 \pm 0.01$ | $0.73 \pm 0.01$ | $0.63 \pm 0.01$ |
| HSP82_YEAST_Bolon2016 | $0.70 \pm 0.01$ | $0.64 \pm 0.02$ | $0.72 \pm 0.01$ | $0.65 \pm 0.02$ |
| KKA2_KLEPN_Mikkelsen2014 | $0.78 \pm 0.02$ | $0.65 \pm 0.04$ | $0.80 \pm 0.01$ | $0.67 \pm 0.03$ |
| MTH3_HAEAESTABILIZED_Tawfik2015 | $0.53 \pm 0.04$ | $0.43 \pm 0.04$ | $0.60 \pm 0.04$ | $0.48 \pm 0.02$ |
| PABP_YEAST_Fields2013-singles | $0.87 \pm 0.02$ | $0.78 \pm 0.03$ | $0.91 \pm 0.01$ | $0.81 \pm 0.03$ |
| POLG_HCVJF_Sun2014 | $0.79 \pm 0.02$ | $0.67 \pm 0.03$ | $0.81 \pm 0.01$ | $0.70 \pm 0.03$ |
| RL401_YEAST_Bolon2013 | $0.82 \pm 0.03$ | $0.71 \pm 0.04$ | $0.84 \pm 0.02$ | $0.72 \pm 0.05$ |
| RL401_YEAST_Bolon2014 | $0.73 \pm 0.02$ | $0.66 \pm 0.05$ | $0.76 \pm 0.02$ | $0.68 \pm 0.05$ |
| UBE4B_MOUSE_Klevit2013-singles | $0.60 \pm 0.06$ | $0.36 \pm 0.09$ | $0.73 \pm 0.06$ | $0.41 \pm 0.08$ |
| YAP1_HUMAN_Fields2012-singles | $0.85 \pm 0.04$ | $0.51 \pm 0.15$ | $0.88 \pm 0.03$ | $0.58 \pm 0.11$ |
